# Degradation pathways for organic matter of terrestrial origin are widespread and expressed in Arctic Ocean microbiomes

**DOI:** 10.1101/2022.08.13.503825

**Authors:** Thomas Grevesse, Céline Guéguen, Vera E. Onana, David A. Walsh

## Abstract

**Background:** The Arctic Ocean receives massive freshwater input and a correspondingly large amount of humic-rich organic matter of terrestrial origin. Global warming, permafrost melt, and a changing hydrological cycle will contribute to an intensification of terrestrial organic matter release to the Arctic Ocean. Although considered recalcitrant to degradation due to complex aromatic structures, humic substances can serve as substrate for microbial growth in terrestrial environments. However, the capacity of marine microbiomes to process aromatic-rich humic substances, and how this processing may contribute to carbon and nutrient cycling in a changing Arctic Ocean, is relatively unexplored. Here, we used a combination of metagenomics and metatranscriptomics to assess the prevalence and diversity of metabolic pathways and bacterial taxa involved in aromatic compound degradation in the salinity-stratified summer waters of the Canada Basin in the western Arctic Ocean.

**Results:** Community-scale meta-omics profiling revealed that 22 complete pathways for processing aromatic compounds were present and expressed in the Canada Basin, including those for aromatic ring fission and upstream funnelling pathways to access diverse aromatic compounds of terrestrial origin. A phylogenetically diverse set of functional marker genes and transcripts were associated with fluorescent dissolved organic matter, a component of which is of terrestrial origin. Pathways were common throughout global ocean microbiomes, but were more abundant in the Canada Basin. Genome-resolved analyses identified 12 clades of *Alphaproteobacteria*, including *Rhodospirillales*, as central contributors to aromatic compound processing. These genomes were mostly restricted in their biogeographical distribution to the Arctic Ocean, and were enriched in aromatic compound processing genes compared to their closest relatives from other oceans.

**Conclusion:** Overall, the detection of a phylogenetically diverse set of genes and transcripts implicated in aromatic compound processing supports the view that Arctic Ocean microbiomes have the capacity to metabolize humic substances of terrestrial origin. In addition, the demonstration that bacterial genomes replete with aromatic compound degradation genes exhibit a limited distribution outside of the Arctic Ocean suggests that processing humic substances is an adaptive trait of the Arctic Ocean microbiome. Future increases in terrestrial organic matter input to the Arctic Ocean may increase the prominence of aromatic compound processing bacteria and their contribution to Arctic carbon and nutrient cycles.

## Introduction

Humic substances (HS) are a heterogeneous mixture of organic compounds resulting from biochemical transformations of dead plants and microbes. HS are ubiquitous in both terrestrial and aquatic systems and constitute the largest fraction of organic matter (OM) in terrestrial ecosystems (Kida *et al*., 2018; Dou *et al*., 2020), reaching 60-80% in soils (Park *et al*., 2015) and 50-80% in freshwaters (Kisand *et al*., 2013). The fraction of HS is relatively high (20-60%) in shelves, coastal zones and estuaries (Esham, Ye and Moran, 2000) due to the input of terrestrial OM (tOM) with freshwater runoff and exchange with sediments. HS constitute a smaller fraction of dissolved OM (DOM) in the open ocean (0.7-2.4%) (Opsahl and Benner, 1997). The lesser amount of HS in the DOM of open oceans indicates that HS is removed by ocean microbiomes and additional non-biological processes (Kisand, Rocker and Simon, 2008; Rocker, Kisand, *et al*., 2012).

The Arctic Ocean receives a disproportionately high input of freshwater (10% of total global freshwater input for 1.3% of total ocean volume), and a correspondingly high tOM input (10% of ocean total tOM input) (Shen *et al*., 2016). Rivers annually discharge 25-36 Tg of dissolved organic carbon and 12 Tg of particulate organic carbon to the Arctic Ocean (Raymond *et al*., 2007; Holmes *et al*., 2012). Climate change is strongly influencing the Arctic region, which in turn is influencing Arctic hydrology and organic matter dynamics (Duarte *et al*., 2012; Macdonald, Kuzyk and Johannessen, 2015). More specifically, permafrost thawing (Frey and McClelland, 2009), combined with intensifying river runoff (Mann *et al*., 2016), coastal erosion (Anderson and Macdonald, 2015) and groundwater input (Connolly *et al*., 2020), is driving an increase in the amount of humic-rich DOM input into the Arctic Ocean. The humic-rich DOM consequently contributes significantly to the carbon pool of the Arctic Ocean DOM compared to other oceans (Gonçalves-Araujo *et al*., 2016), and potentially represents a significant and increasing growth resource for the Arctic Ocean microbiome.

The origins and distributions of tOM in the Canada Basin in the western Arctic Ocean has been extensively studied, making it a useful system for investigating interactions between tOM and ocean microbiomes. In spring and summer, humic-rich OM is transported by riverine inputs to the surface mixed layer of the Arctic Ocean shelves (Muscolo *et al*., 2013; Polyakov *et al*., 2018). In shelf waters, tOM is partially photodegraded (Chupakova *et al*., 2018), while some flocculates upon mixing with salt water and sinks to the sediments along with particulate OM (Mann *et al*., 2016). In fall and winter, the tOM remaining in the surface layer sinks with the dense brine expelled during ice-formation. This brine flows along Chukchi Sea and Beaufort Sea shelves, exchanging organic matter with bottom sediments, ultimately accumulating in the deeper and more saline water of Pacific Ocean origin (Anderson and Macdonald, 2015; Jung *et al*., 2021). The interactions with shelf sediments and pore waters constitute a substantial source of tOM which may have been reprocessed by sediment microbiomes (Chen *et al*., 2016). It has been estimated that 11-44% of Arctic Ocean sediment OM is of terrestrial origin (Belicka and Harvey, 2009). As a consequence of the OM dynamics, the Canada Basin is characterized by a strong and distinctive signal of humic-rich DOM that extends from the subsurface water to a depth of ∼300 m (Gao and Guéguen, 2018; DeFrancesco and Guéguen, 2021).

HS are heterogeneous supramolecular assemblies formed by microbial and physico-chemical transformations (Sutton and Sposito, 2005) of organic matter. In terrestrial systems, HS originates from vascular plant residues (lignin and other biopolymers) and other organic detritus (Gerke, 2018), giving rise to HS rich in aromatic moieties. In contrast, HS produced in marine environments have a strong aliphatic and branched structure (Esteves, Otero and Duarte, 2009). In the Arctic Ocean however, HS are aromatic-rich due to their terrestrial and sediment origin (Dittmar and Kattner, 2003). HS usually show a high degree of recalcitrance that is dependent on their physicochemical interactions with the environment (Hedges *et al*., 2000). In soils, sorption of HS to mineral particles drives a physical separation of HS from microbes and their enzymes, preventing fast degradation of HS (Ekschmitt *et al*., 2008; Schmidt *et al*., 2011). Numerous studies have therefore demonstrated that HS freed from their soil environment can be used to support microbial growth (Hertkorn *et al*., 2002; Kisand, Rocker and Simon, 2008; Kim *et al*., 2019).

The capacity of microbiomes to couple HS transformation to growth relies on the ability to degrade aromatic compounds from HS. The degradation of aromatic compounds follows two main steps. Funneling pathways transform (*e*.*g*. via oxidation, decarboxylation, and/or demethylation) larger and more substituted aromatic compounds to a small set of key aromatic compounds (*e*.*g*. gentisate, catechol, protocatechuate), which then undergo an aromatic ring-fission step followed by further processing to generate central carbon metabolism intermediates. In humic-rich environments such as soils, microbiomes use a wide variety of funneling pathways to access the diverse set of lignin-derived aromatic compounds (*e*.*g*. vanillate, syringate, benzoate and their derivatives) that have been incorporated in HS (Brink *et al*., 2019).

In soils, fungi degrade most of the humic substances (Janusz *et al*., 2017). In the ocean, bacteria are considered the main actors in OM degradation (Herndl *et al*., 2008), even if recent studies have highlighted an important role for fungi (Comeau *et al*., 2016; Gladfelter, James and Amend, 2019), for example in processing OM in marine snow (Bochdansky, Clouse and Herndl, 2017) or by parasitizing phytoplankton (Ilicic and Grossart, 2022). Certain bacteria inhabiting humic-rich environments can grow on HS as sole carbon and energy sources, and are therefore are able to access aromatic compounds within HS (Esham, Ye and Moran, 2000; Rocker, Brinkhoff, *et al*., 2012; Kim *et al*., 2019). Transcriptomic analysis from the humic acid-degrading bacterium *Pseudomonas sp*. isolated from sub-arctic tundra soils showed that genes involved in the funneling and ring-opening steps of AC degradation pathways were up-regulated when fed with humic acids compared to glucose (Kim *et al*., 2018). Recently it was shown that *Chloroflexi* genomes from the Canada Basin encoded a diverse set of genes associated with aromatic compound degradation (Colatriano *et al*., 2018). The *Chloroflexi* populations appeared to be endemic to the Arctic Ocean and were associated with the humic-rich fluorescence DOM maximum (FDOMmax). These observations suggest the disproportionately high fraction and diversity of aromatic-rich HS in the Arctic Ocean DOM compared to other oceans may select for a diverse HS-degrading microbiome.

The genomic diversity and metabolic pathways in the Arctic Ocean microbiomes can provide important insights regarding the fate of HS and its impact on Arctic Ocean biogeochemical cycles. However, outside of perhaps the *Chloroflexi*, we know very little about how phylogenetically widespread HS degradation is in the Arctic Ocean microbiomes, nor the diversity of metabolic pathways employed by the Arctic Ocean microbiomes to process HS. We hypothesized that the capacity for aromatic compound degradation was linked to the distribution of humic-rich tOM and enhanced in the Arctic Ocean compared to other oceans with a lesser amount of HS. Finally, we hypothesized that the vast amount of HS in the Arctic Ocean may have played a role as an ecological pressure for the adaptive evolution of the taxa most implicated in aromatic compound degradation.

## Results

### Environmental context

We surveyed the microbiomes along a latitudinal transect (73^-^81 °N) of the salinity-stratified waters of the Canada Basin using a combination of metagenomics and metatranscriptomics (**Figure 1a-b, Table S1**). The sampling design targeted distinct water column features, including the relatively fresh surface mixed layer (surface; 5 m and 20 m), the subsurface chlorophyll maximum (SCM; 55-95 m), the FDOMmax associated with colder Pacific-origin water (32.3 and 33.1 PSU; 90-250 m), the warmer Atlantic-origin water (Tmax and AW; 360-1000 m depth), and Arctic bottom water (∼3800 m). The warmer Atlantic-origin water and Arctic bottom water are herein collectively referred to as deep waters.

**Figure 1.**
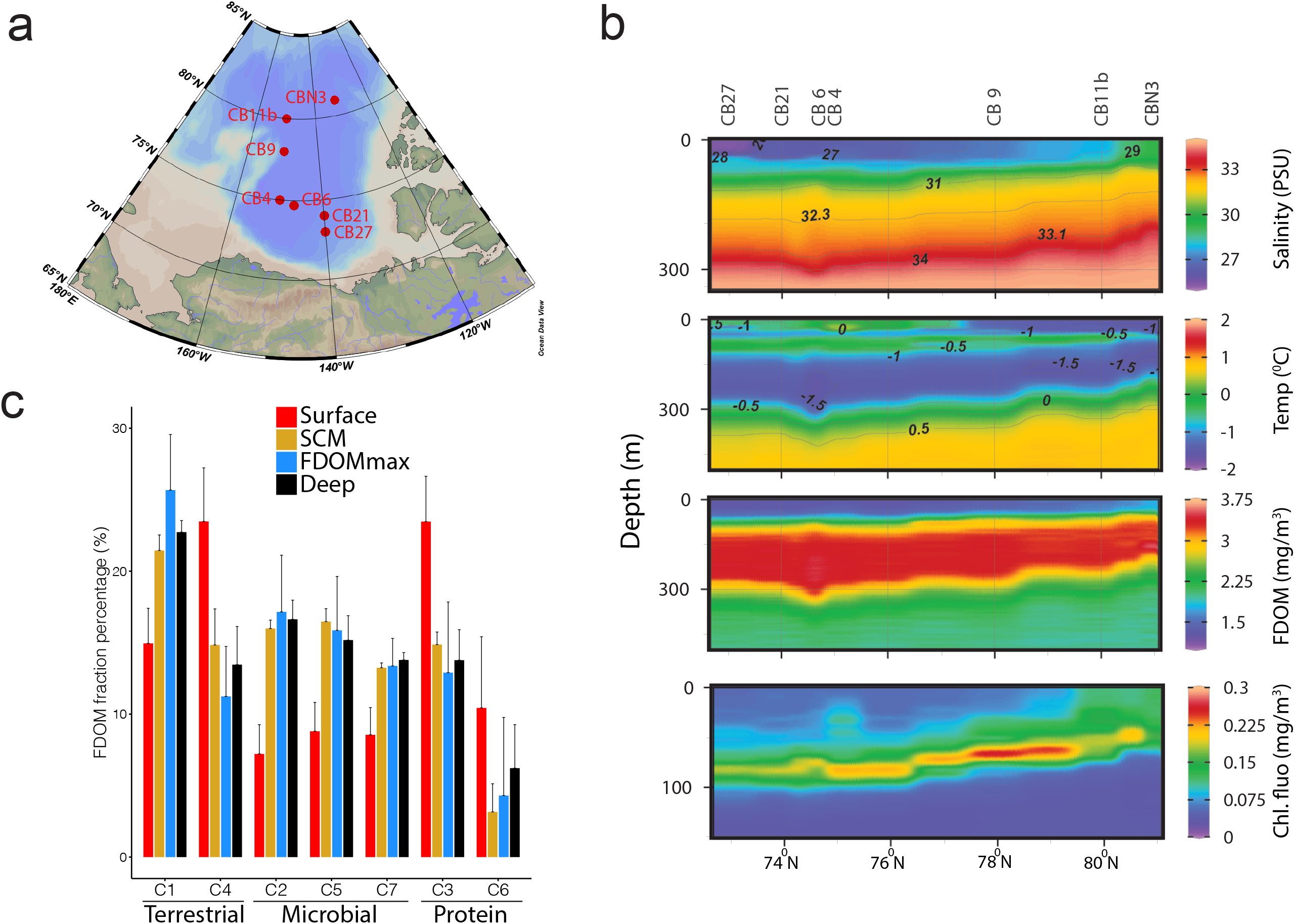
Spatial biogeochemistry of the Canada Basin. a) Map of the 7 stations sampled in this study. b) depth profile of Salinity (PSU), temperature (^0^C), fluorescent dissolved organic matter (FDOM, mg/m^3^), and chlorophyll fluorescence (mg/m^3^) at the 8 stations sampled in this study. c) Percentage of the 7 FDOM fractions identified using excitation emission matrix fluorescence spectroscopy combined with parallel factor analysis. Samples are grouped in 4 samples water features: Surface, subsurface chlorophyll maximum (SCM) fluorescent dissolved organic matter maximum (FDOMmax) and deep waters.

We sought to determine the distribution and composition of OM in the Canada Basin, with a focus on the distribution of tOM. Optical properties of the OM, such as fluorescence, have previously been used to assess the composition of OM in the ocean, and differentiate between terrestrial and marine OM sources (Murphy *et al*., 2008; Guéguen and Kowalczuk, 2013). We used excitation emission matrix (EEC) fluorescence spectroscopy combined with parallel factor analysis (PARAFAC) to determine the distribution of fluorescent DOM components. In the Canada Basin, seven components (C1-C7) were identified, as previously defined in (DeFrancesco and Guéguen, 2021). These components corresponded to terrestrially derived humic-like DOM (C1 and C4), amino acid or protein material (C3 and C6), or microbially-derived humic-like DOM (C2, C5 and C7) (**Figure 1c**). The aromatic-rich C1 was the most abundant component within the FDOMmax samples (25-27 %), but also in the whole water column below the surface (20-22% in the SCM and 21-23% in the deep), verifying that a significant fraction of OM is of terrestrial origin. Of the terrestrial components, C4 was the dominant component in the surface (19-30 %). The reduced contribution of C1 in the surface is because C1 is more red-shifted than C4 indicating a stronger aromatic character and thus enhanced photosensitivity. Overall, these results indicate a strong contribution of a photostable fraction from terrestrial origin in the FDOM of the surface and an aromatic-rich fraction from terrestrial origin in the FDOM of the whole water column below the surface.

### Vertical-partitioning of metabolic features in metagenomes and metatranscriptomes

We investigated the abundance and distribution of aromatic compound degradation pathways in the Canada Basin microbiomes in relation to tOM availability. To investigate how metabolic pathways of were distributed across the Canada Basin, we first performed nonmetric multidimensional scaling (NMDS) analysis on the abundance of enzyme-encoding genes (genes assigned to enzyme commission (EC) numbers) annotated from metagenome or metatranscriptome assemblies. NMDS ordination showed that metagenomes (stress=0.11) were partitioned into four clusters consisting of samples collected from either the surface, the SCM, FDOMmax, or deep water (**Figure 2a**). A similar pattern was observed in the NMDS ordination of metatranscriptomes (stress=0.0503), although the variation between samples from within the same water column feature was higher than observed in metagenomes (**Figure 2b**). In addition, there was less separation between the samples from the FDOMmax and from deeper Atlantic-origin waters in the ordination based on metatranscriptomes compared to metagenomes.

**Figure 2.**
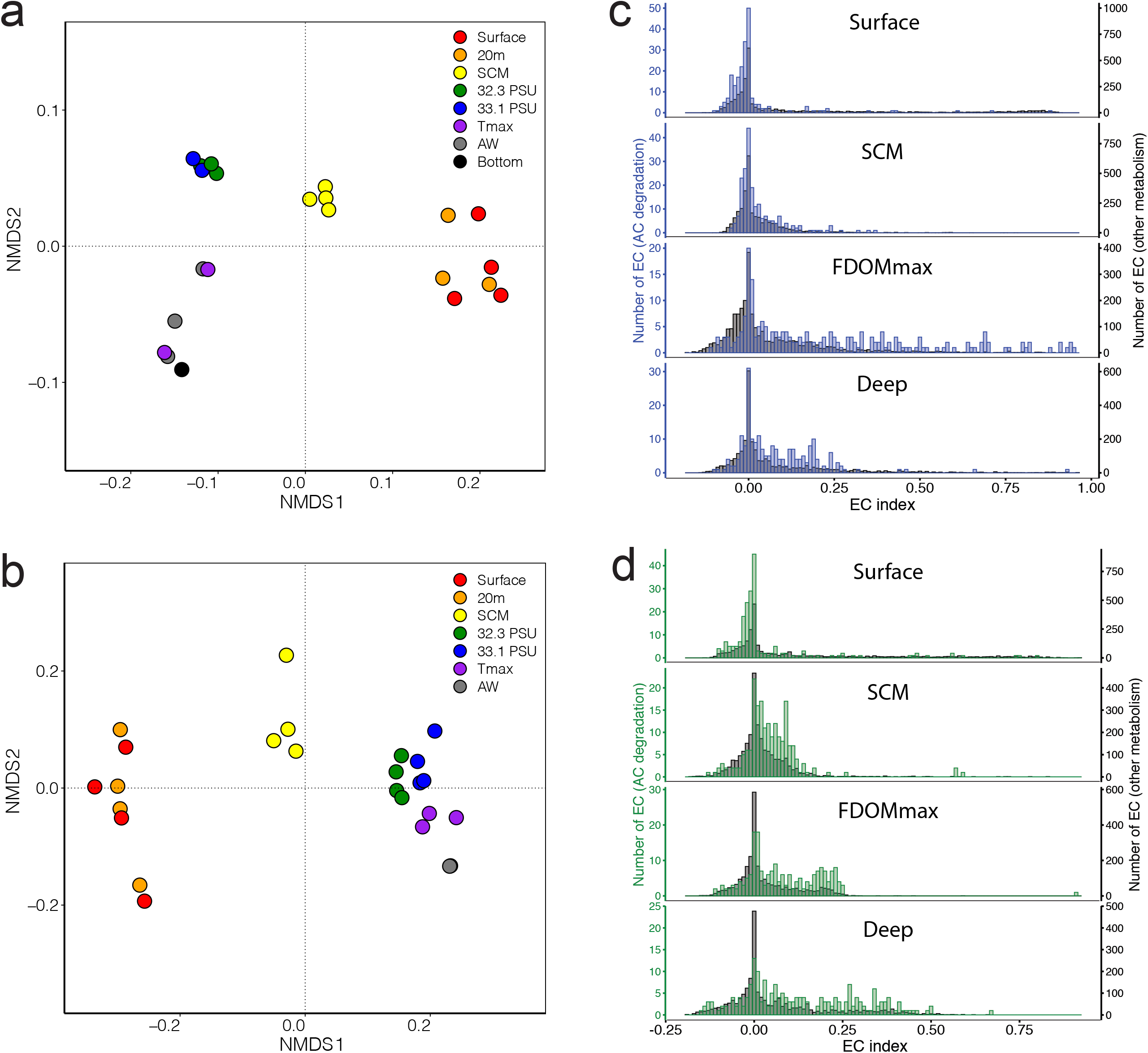
Identification of the microbial enzymatic reactions associated with the 4 water column features. Non metric multidimensional scaling of the matrices of enzymatic reactions abundances from a) metagenomes and b) metatranscriptomes. Distribution of gene indices calculated for the enzymatic reactions from c) metagenomes and d) metatranscriptomes in the 4 water features. Blue/green: distribution for enzymatic reactions involved in the degradation of aromatic compounds from metagenomes (blue) and metatranscriptomes (green); grey: distribution for all enzymatic reactions not involved in the degradation of aromatic compounds.

We next determined which enzymatic reactions differentiated the metagenomes across the stratified water column using non-negative matrix factorization (NMF), which is a tool for extracting meaningful features from high dimensional data (Seung and Lee, 1999). In our analyses, NMF decomposes the matrix of EC number abundances into two matrices. Matrix 1 presents a reduced number of elements that describe the overall similarities of the metagenomes based on EC number composition, while matrix 2 presents the weighted contribution of individual EC numbers on each of the elements in matrix 1. We determined that a decomposition with four elements best represented the overall enzyme composition of metagenomes (**Figure S1)**. The four elements, herein referred to as sub-metagenomes (**Figure S2**), represented the same patterns as observed in the NMDS ordination, corresponding to the surface, SCM, FDOMmax, and deep waters (**Figure 2a**).

We then assessed which EC numbers were strongly associated with each of the four sub-metagenomes by calculating an EC index value. This EC index value quantifies the tendency of an EC number to be specific to a single sub-metagenome (EC index values range between -1 and 1). The distribution of EC indices was plotted for each of the four sub-metagenomes. Overall, the means of the EC indices associated with aromatic compound degradation and other metabolic pathways in the four water column features were significantly different (PERMANOVA, F=89.8, p<0.0001). Each sub-metagenome has a collection of EC numbers with relatively high indices (>0.5) (**Figure 2c**). However, the most striking observation was that EC numbers involved in aromatic compound degradation were predominantly associated with the FDOMmax sub-metagenome, as demonstrated by the higher index values for EC numbers from aromatic compound degradation pathways than from other metabolic pathways in the FDOMmax (Student t-test, t=13.26, p<0.0001). The EC indices for aromatic compound degradation genes were smaller than the EC indices associated with other metabolic processes in the surface (Student t-test, t=8.89, p<0.0001) and not significantly different for the SCM (Student t-test, t=0.369, p=0.414) and the deep (Student t-test, t=0.56, p=0.545) sub-metagenomes.

We performed a similar NMF analysis on EC numbers in the metatranscriptomes (**Figure S1**). Similar with the NMF analysis of metagenomes, decomposition resulted in four elements, herein referred to as sub-metatranscriptomes (**Figure S1**), corresponding to the surface, SCM, FDOMmax, and deep waters (**Figure S3)**. For the sub-metatranscriptomes, the means of the EC indices from aromatic compound degradation and from other metabolic pathways in the four water column features were significantly different (PERMANOVA, F=121, p<0.0001). For the sub-metatranscriptomes, however, we observed higher indices for the EC numbers from aromatic compound degradation than for EC numbers from other metabolic processes in the SCM (Student t-test, t=0.0218, p<0.0001), the FDOMmax (Student t-test, t=0.0444, p<0.0001) and the deep waters (Student t-test, t=0.0611, p<0.0001) (**Figure 2d**).

### Aromatic compound degradation genes in global ocean metagenomes

As the humic-rich OM input to the Arctic Ocean is disproportionately high compared to other oceans, we investigated if genes associated with aromatic compound degradation were more abundant in the Canada Basin metagenomes compared to other oceanic metagenomes. As terrestrial OM is a significant contributor of HS to the Arctic Ocean, we restricted our analysis to genes involved in processing aromatic compounds of terrestrial origin. We focused the analysis on a set of 46 pathways previously implicated in degrading aromatic compounds from lignin (**Figure 3a, Table S2**). We compared the relative abundance of aromatic compound degradation genes between metagenomes of the Canada Basin water features (surface, SCM, FDOMmax, deep) and metagenomes from both the surface and subsurface waters (SCM, mesopelagic, bathypelagic) of the Atlantic, Pacific, Indian and Southern Oceans as well as the Mediterranean Sea and Red Sea (**Figure 3b**). The Canada Basin FDOMmax metagenomes contained the highest fraction of aromatic compound degradation genes (3.4%). Aromatic compound degradation genes were identified in other oceanic metagenomes (1.5-2.5 % of total protein-coding genes) and the relative abundance of aromatic compound degradation genes increased with water depth. Overall, the mean percentage of aromatic compound degradation genes in the water column features of the Canada Basin were significantly different than in other oceans (PERMANOVA, F=27.8, p<0.0001). Specifically, the relative abundance was consistently higher (1.3-1.7 fold) in the microbiomes of the Canada Basin upper water column features compared to microbiomes from other oceanic zones (Student t-test: surface/surface, t=0.58, p=0.0013; SCM/SCM, t=0.92, p=0.0001; FDOMmax/mesopelagic, t=0.81, p=0.0001) (**Figure 3c**). However, we did not observe significant differences between the percentage of aromatic compound degradation genes of Arctic deep-water microbiomes and the microbiomes of other oceans deep waters (Student t-test, t=0.20, p=0.490) (**Figure 3c**).

**Figure 3.**
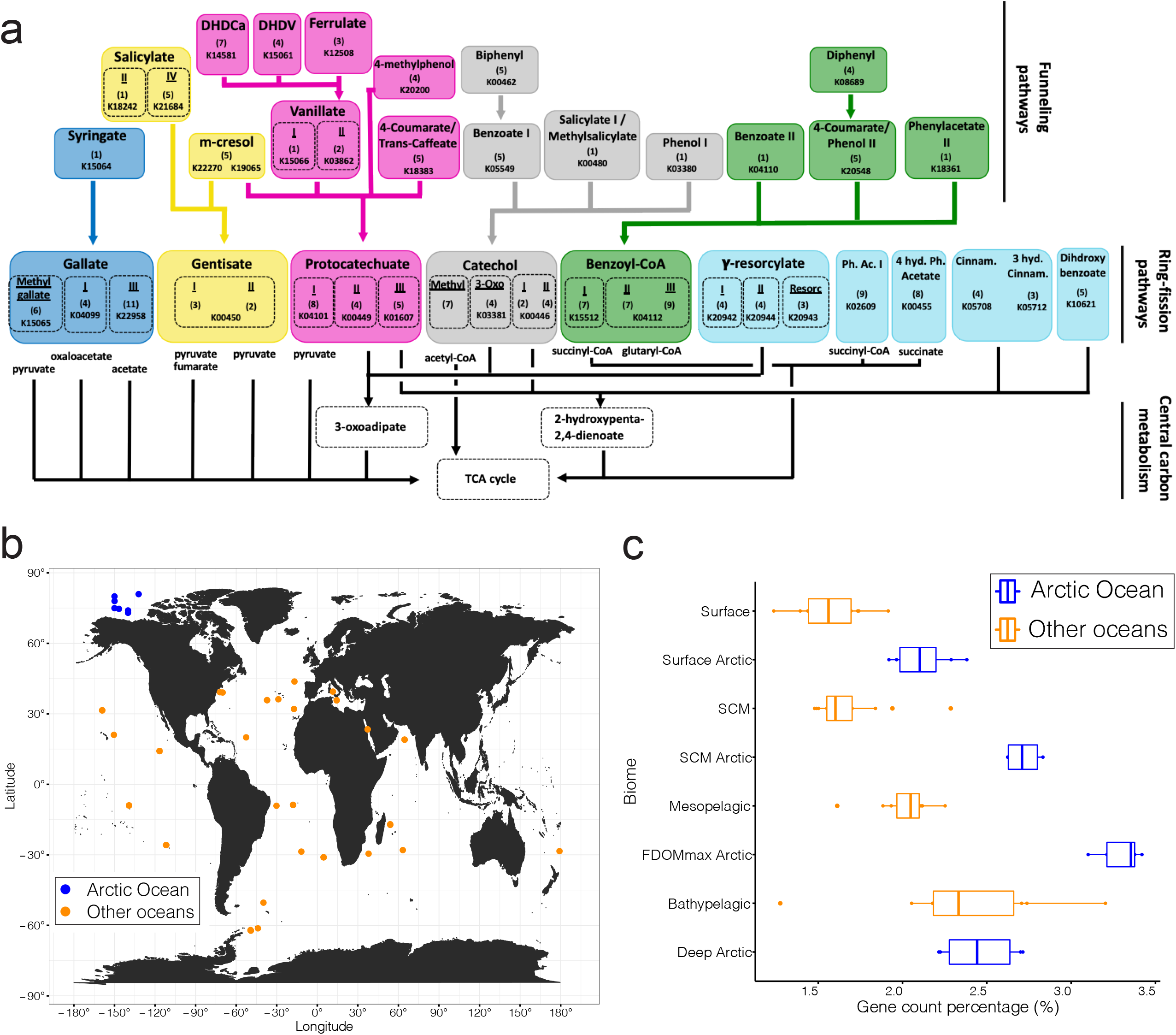
a) Diagram of the 46 funneling and ring-opening pathways involved in the degradation of lignin-derived aromatic compounds. b) Map of the metagenomic samples used to compare the capacity to degrade aromatic compounds in the Arctic Ocean (blue) compared to other seas and oceans (orange). c) Fraction of metabolic genes involved in the degradation of aromatic compounds in the Arctic Ocean (blue) and other seas and oceans (orange). The line in the box represents the median. The left and right hinges of the box represent the 25^th^ and 75^th^ percentiles. Whiskers extend to the smallest and largest values (no further than 1.5* the inter-quartile range)

### Distribution of aromatic compound degradation genes and pathways in metagenomes and metatranscriptomes

To elucidate the diversity of aromatic compounds that the Arctic Ocean microbiomes can access as growth substrates, we assessed the diversity and the completeness of aromatic compound degradation pathways in Canada Basin metagenomes. We found evidence for the presence of 44 of the 46 aromatic compound degradation pathways in the metagenomes (**Figure S4**). A complete set of genes were identified for over half of these pathways in the metagenomes, irrespective of the water column feature (**Figure S4**). Evidence for the 44 pathways was also identified in the metatranscriptomes, including expression of the full complement of genes for 22 pathways (**Figure S4**).

To measure the distribution of the aromatic compound degradation pathways through the water column, we used a selection of 39 unique marker genes for the 46 aromatic compound degradation pathways (**Table S2**). To provide a measure of pathway abundance and expression, we summed the depth of coverage of each marker gene or transcript and corrected for differences in metagenome sequencing effort (**Figure 4**). Out of the 39 unique marker genes, 32 were detected in the Canada Basin metagenomes (**Figure 4a, Figure S5**). Generally, the most abundant genes were also most abundant in the metatranscriptomes (**Figure 4a-b, Figure S5, Figure S6**). Most of the marker genes were most abundant in the FDOMmax, yet were most highly expressed in deep waters. (**Figure 4a-b**).

**Figure 4.**
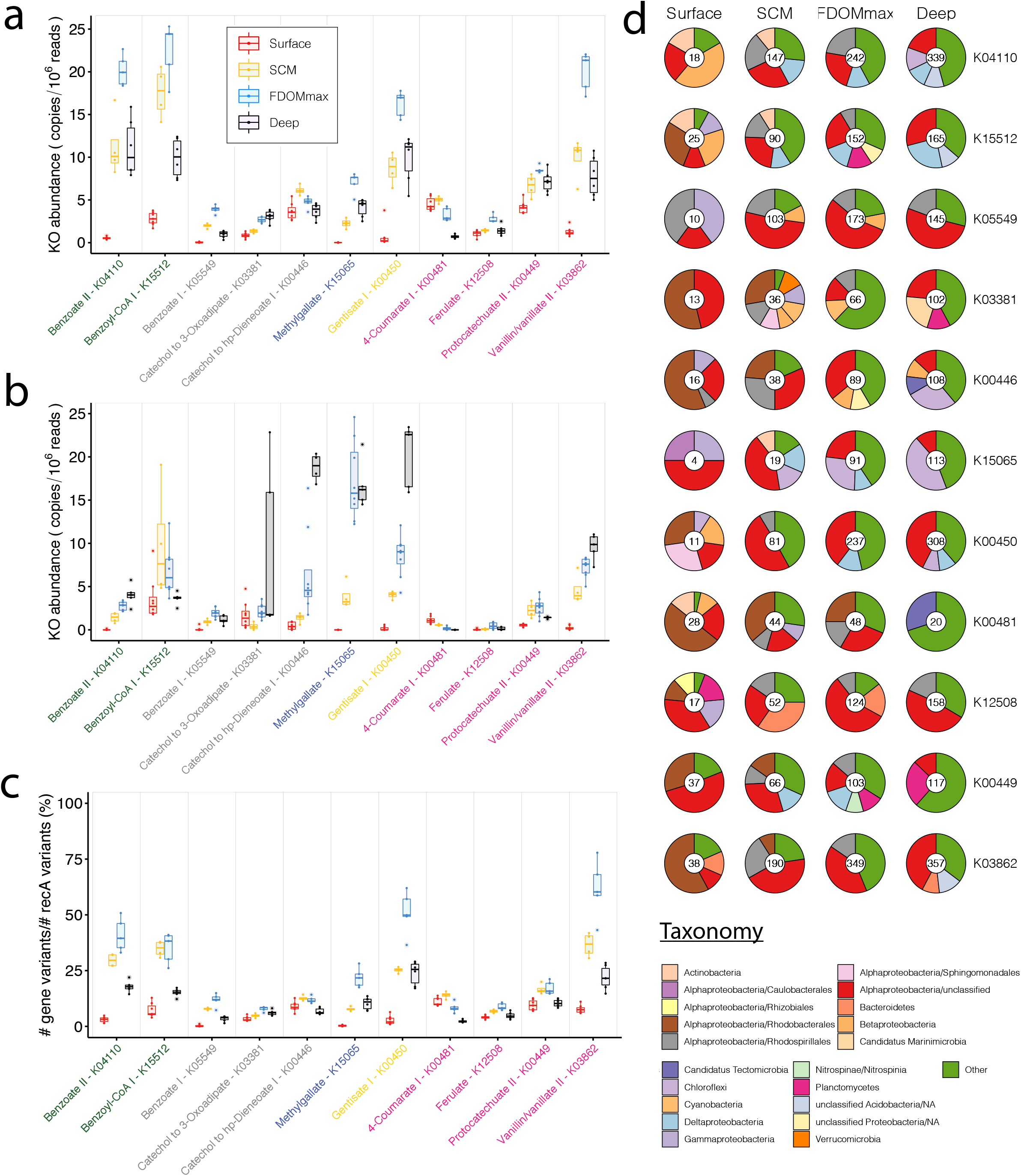
Distribution and taxonomy of marker genes involved in the degradation of aromatic compounds in the Arctic Ocean. a) normalized abundance of genes annotated with KEGG orthology numbers (KOs) marker of aromatic degradation pathways in the 4 different water features. b) normalized abundance of transcripts annotated with KEGG orthology numbers marker of aromatic degradation pathways in the 4 different water features. c) estimated fraction of the microbiomes harboring genes annotated with KOs marker of aromatic degradation pathways in the 4 different water features. The line in boxes represents the medians. The bottom and upper hinges of the box represent the 25^th^ and 75^th^ percentiles. Whiskers extend to the smallest and largest values (no further than 1.5* the inter-quartile range). d) taxonomy of genes annotated with KOs marker of aromatic degradation pathways in the 4 different water features. Numbers within pie charts represent the number of gene clusters (clustered at 95% identity) identified in each water feature. Red: surface samples; yellow: SCM samples; blue: FDOMmax samples; black: deep samples.

Vanillate monooxygenase (K03862) was the most abundant marker gene within all water column features (20 copies/10^6^ reads in the FDOMmax) for pathways degrading aromatic compounds from terrestrial sources, while 3-O-methylgallate 3,4-dioxygenase (K15065) showed a lower abundance (8 copies/10^6^ reads in the FDOMmax). While vanillate monooxygenase was more abundant in the metatranscriptomes of the FDOMmax, 3-O-methylgallate 3,4-dioxygenase was more abundant in the metatranscriptomes of the deep layers. Both ring fission protocatechuate dioxygenases (K00449 and K04101) were abundant in metagenomes (8 and 6 copies/10^6^ reads in the FDOMmax) and metatranscriptomes (2.5 and 7.5 copies/10^6^ reads in the FDOMmax). Overall, these results show that the Canada Basin microbiomes can fully transform aromatic compounds from terrestrial sources into central carbon metabolism intermediates, with an enhanced capacity in the FDOMmax.

A number of aromatic compounds (*e*.*g*. salicylate, 3-hydroxycinnamate or benzoate) can originate from lignin as well as other sources such as marine phytoplankton. Within the pathways involved in the degradation of aromatic compounds from possible marine or terrestrial origin, benzoate CoA-ligase (K04110), salicylate monooxygenase (K00480) and 3-hydroxycinnamate hydroxylase (K05712) were the most abundant in metagenomes (20, 24 and 17 copies/10^6^ reads respectively) but not in metatranscriptomes (7.5, 5 and 2 copies/10^6^ reads). The most common funneling pathway for benzoate was through benzoyl-CoA as evidenced by the lower abundance of genes (5 copies/10^6^ reads) and transcripts (2 copies/10^6^ reads) encoding benzoate 1,2-dioxygenase (K05549) compared to benzoate CoA-ligase. Accordingly, the ring-fission benzoyl-CoA 2,3-epoxidase (K15512) was significantly more abundant (22 copies/10^6^ reads) than the ring-fission marker genes catechol 1,2-dioxygenase (K03381) and catechol 2,3-dioxygenase (K00446) (3 and 7 copies/10^6^ reads in the deep and SCM respectively). However, both benzoyl-CoA 2,3-epoxidase and catechol 2,3-dioxygenase were among the most abundant genes in the metatranscriptomes (20 copies/10^6^ reads), but with maximum abundance in the SCM and the deep, respectively (**Figure 4b**). Of the ring-fission pathway marker genes, gentisate 1,2-dioxygenase (K00450) was one of the most abundant in metagenomes (15 copies/10^6^ reads in the FDOMmax) and metatranscriptomes (20 copies/10^6^ reads in the deep).

### Taxonomic identity of aromatic compound degradation genes and their distribution across the microbiomes

We estimated the fraction of bacterial genomes harboring each marker gene by comparing the total number of gene variants for select aromatic compound degradation pathway markers to the number of the single copy universally-distributed *recA* genes (**Figure 4c, Figure S7**). The estimated fraction of bacterial genomes with AC degradation genes increased with depth, reaching a maximum in the FDOMmax (8-75%, **Figure 4c**) and then decreased in the deep water (5-25%). The genes present in the highest fraction of bacterial genomes were involved in the degradation of benzoate through benzoyl-CoA (50% for K04110 and 45% for K15521 in the FDOMmax), gentisate (65% for K00450 in the FDOMmax), vanillate (75% for K03862 in the FDOMmax), salicylate and 3-hydroxycinnamate (45% for K00480 and 40% for K05712 in the SCM) (**Figure S7**). These numbers may be overestimated as they assume only a single gene copy per genome, whereas multiple paralogs may be present in a single genome (continued below).

Taxonomic analysis of aromatic compound degradation marker genes revealed that the number of gene clusters generally increased continuously with depth (**Figure 4d**). Surface gene clusters were predominantly affiliated with *Rhodobacterales* (more than 50% of the gene clusters for K03381, K00446, K00481 and K03862) and unclassified *Alphaproteobacteria* (up to 50% for K15065 gene clusters), with a significant contribution from *Gammaproteobacteria* for K05549 (40%) and K15065 (25%) (**Figure 4d**). In the SCM and FDOMmax, unclassified *Alphaproteobacteria* dominated the taxonomic affiliations of aromatic compound degradation gene clusters (10-55%) and *Rhodospirillales* contributed significantly to all gene clusters (10-30%), except for genes involved in the degradation of methylgallate (K15065), which was primarily encoded by *Chloroflexi* (30% in the FDOMmax) (**Figure 4d**). We generally observed more gene clusters in the deep than in the FDOMmax (**Figure 4d**), while these genes were present in a smaller fraction of the deep communities than the FDOMmax communities (**Figure 4c**), suggesting a broader phylogenetic diversity of aromatic compound degradation genes in the deep than in the FDOMmax. This is supported by the large contribution of other taxa (taxa contributing individually to <5%) in the deep microbiomes. The contribution of taxa such as *Rhodospirillales* and *Rhodobacterales* may be underestimated due to the large fraction of *Alphaproteobacteria* genes that could only be assigned at the class level.

### Aromatic compounds processing pathways captured in metagenome assembled genomes

We reconstructed metagenome-assembled genomes (MAGs) from our metagenomic data. We performed metagenomic binning of each of the 22 metagenome assemblies individually to reconstruct a total of 1,772 MAGs. After filtering for genomes with greater than 30% completeness and less than 10% contamination, 823 genomes remained (**Figure S8)**. Thirty-one of the 32 marker genes involved in aromatic compound degradation pathways were identified (only dihydroxyphenylacetate 2,3-dioxygenase – K00455 was not detected) across 59% (482 of 823) of the MAGs (**Figure S9 and S10**). The highest percentage of MAGs harboring aromatic compound degradation genes was in the FDOMmax (64%) and SCM (67%), and the lowest percentage in the surface (47%) and deep waters (54%). In general, the taxonomic diversity of MAGs increased with depth. Marker genes were identified in a broad taxonomic diversity of MAGs, including *Alphaproteobacteria, Gammaproteobacteria*, and *Dehalococcoidiia*, among others (**Figure S9**). *Alphaproteobacteria* were common in the SCM and FDOMmax, while *Gammaproteobacteria* were common in the surface.

To investigate the ecology of bacterial taxa most implicated in the degradation of aromatic compounds, we further examined MAGs with complete or near-complete aromatic compound degradation pathways. We selected 46 MAGs most enriched in near-complete aromatic compound degradation pathways (see methods, **Figure S10, Table S3**). Of the 46 MAGs, 24 were recovered from metagenomes originating from the FDOMmax and 16 from the SCM layers. 38 of the MAGs were assigned to *Alphaproteobacteria* (**Figure S11**), 3 MAGs belonged to the *Dehalococcoida*, 4 MAGs to the *Gammaproteobacteria* and one MAG to the class *Binatia*.

Given the large representation of *Alphaproteobacteria* in the MAGs most implicated in aromatic compounds degradation, we investigated the evolutionary origins and phylogenetic relationships between our 38 *Alphaproteobacteria* MAGs and reference genomes available from the Genome Taxonomy Database (GTDB) (Chaumeil *et al*., 2020) **(Table S4)**. Based on an average nucleotide identity threshold of 95%, our 38 *Alphaproteobacteria* MAGs belonged to 16 genomospecies, that were phylogenetically associated with 12 clades (**Figure 5a, Figure S12**). The clades were within the *Rhodobacterales, Thallassobaculales, Rhodospirillales, Defluviicoccales* and five GTDB orders of uncultured *Alphaproteobacteria* (UBA8366, UBA6615, GCA2731375, UBA2966, UBA828). Each clade was comprised of Canada Basin MAGs as well as a basal branch consisting of genomes of marine origin. These results demonstrate that the *Alphaproteobacteria* MAGs were phylogenetically distinct but shared recent common ancestry with genomes from other oceanic zones.

**Figure 5.**
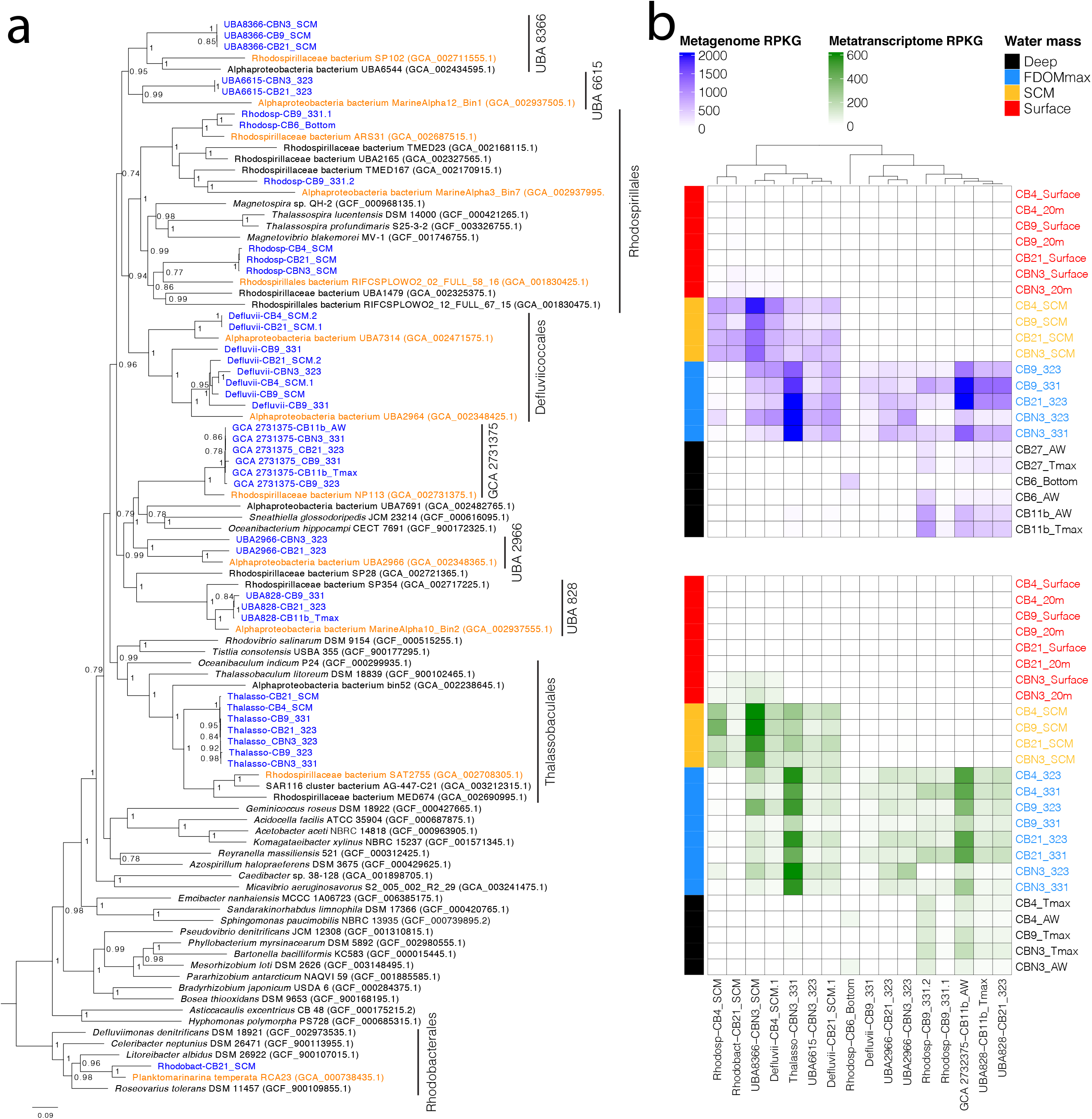
a) Phylogenetic tree of the *Alphaproteobacteria* MAGs identified as most implicated in the degradation of aromatic compounds in the Arctic Ocean. Blue: MAGs reconstructed in this study; orange: genomes from other studies identified as the closest relatives to the MAGs reconstructed in this study; black: other genomes. b) geographic distribution of the *Alphaproteobacteria* MAGs identified as most implicated in the degradation of aromatic compounds within the Canada basin. Top panel: mapping of the metagenomes on the MAGs. Bottom panel: mapping of the metatranscriptomes on the MAGs.

To investigate the distribution of the 12 clades represented by the *Alphaproteobacteria* MAGs across the Arctic water column features, we performed fragment recruitment of both the metagenomic and metatranscriptomic reads against the MAGs representing the 16 genomospecies (**Figure 5b**). Overall, the distribution in metagenomes and metatranscriptomes was similar and all the MAGs were most abundant and active either in the FDOMmax or the SCM (**Figure 5b**). We identified four general patterns of distribution across water column features consisting of 1) restriction to the FDOMmax (Defluvii-CB9_331, UBA2966), 2) common to the SCM and FDOMmax (UBA8366, UBA6615 clade, *Thalassobaculales* clade, *Defluviiccocales* genomes CB21_SCM.1 and CB4_SCM.1), 3) common in the FDOMmax and deeper waters (UBA828 clade genomes, GCA 2732375 genomes and *Rhodospirillales* genomes) and 4) restricted to the SCM (Rhodosp-CB4_SCM and Rhodobact-CB21_SCM). These results show that MAGs with near-complete aromatic compound degradation pathways are strongly associated with and active in HS-rich regions of the water column.

### Global ocean distribution of Alphaproteobacteria MAGs from Canada Basin

We sought to determine if the MAGs most implicated in aromatic compound degradation were more broadly distributed beyond the Arctic Ocean. We therefore investigated the distribution of the *Alphaproteobacteria* MAGs (**Table S5**) by fragment recruitment against a set of 151 metagenomes broadly representative of the global ocean microbiome (**Figure 6**). Of the 16 representative MAGs, two were commonly detected outside of the Arctic Ocean. Rhodosp-CB9_331.2 was identified in the mesopelagic metagenomes from all oceanic regions, but not the Mediterranean Sea or Red Sea. The *Rhodobacterales* MAG (Rhodobact-CB21_SCM) was also identified in surface water metagenomes, most notably from the Southern Ocean. Although several other MAGs were detected at low frequency in the Southern Ocean (*e*.*g*. Rhodosp-CB6-bottom and Defluvii-CB9-331) the majority (12 MAGs, 75% of the MAGs) were not detected outside of the Arctic Ocean. Likewise, the vast majority of the most closely related marine reference genomes were not detected in Canada Basin metagenomes. The exceptions were *Planktomarinamarina* (*Rhodobacterales*) and the reference genome within UBA828.

**Figure 6.**
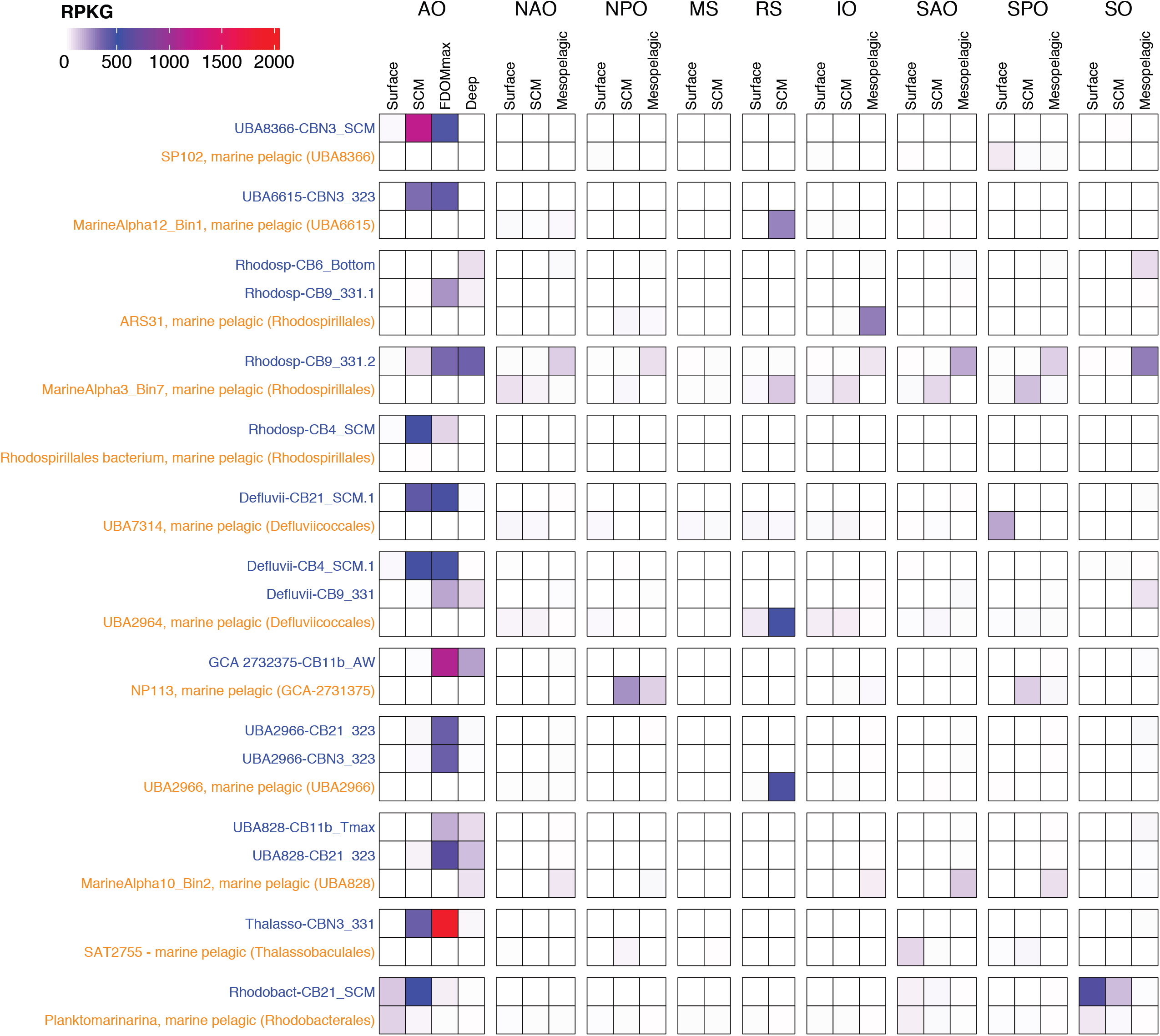
Fragment recruitment of global oceans metagenomes on the *Alphaproteobacteria* MAGs identified as most implicated in the degradation of aromatic compounds. Blue: MAGs identified in this study and representative of the 16 genomospecies; orange: genomes from other studies identified as the closest relatives to the MAGs reconstructed in this study. AO: Arctic Ocean; IO: Indian Ocean; MS: Mediterranean Sea; NAO: North Atlantic Ocean; NPO: North Pacific Ocean; RS: Red Sea; SAO: Southern Atlantic Ocean; SO: Southern Ocean; SPO: Southern Pacific Ocean.

### Enhanced aromatic compound degradation capacity in *Alphaproteobacteria* MAGs restricted to the Canada Basin

We hypothesized that an enhanced capability for aromatic compound degradation may be implicated in the evolutionary adaptation of the Arctic Ocean populations. We therefore compared the abundance and diversity of aromatic compound degradation genes between the Arctic MAGs and the set of their closest relatives. Between 1.5% and 4% of the genes with an EC annotation was annotated with an EC from lignin-derived aromatic compound degradation pathways for both the Arctic MAGs and genomes from other oceans (**Figure 7**). However, out of the 16 Arctic Ocean MAGs, 10 possessed a higher diversity of marker genes and 14 possessed a higher number of marker gene copies than their sister taxa from other oceans (**Figure 7**). This is despite the difference in genome completeness estimated for the reference MAGs (64-98%) compared to the Arctic Ocean MAGs (range of 42-95% completeness). Some marker genes were found exclusively in our Arctic *Alphaproteobacteria* MAGs, including those for the degradation of catechol to 3-oxoadipate, phenol I, dehydrodivanillate, protocatechuate I, cinnamate and resorcinol (**Figure 7**). Other marker genes were found exclusively in the reference MAGs, such as genes for the degradation of benzoyl-CoA II, gallate I and III, m-cresol and γ-resorcylate.

**Figure 7.**
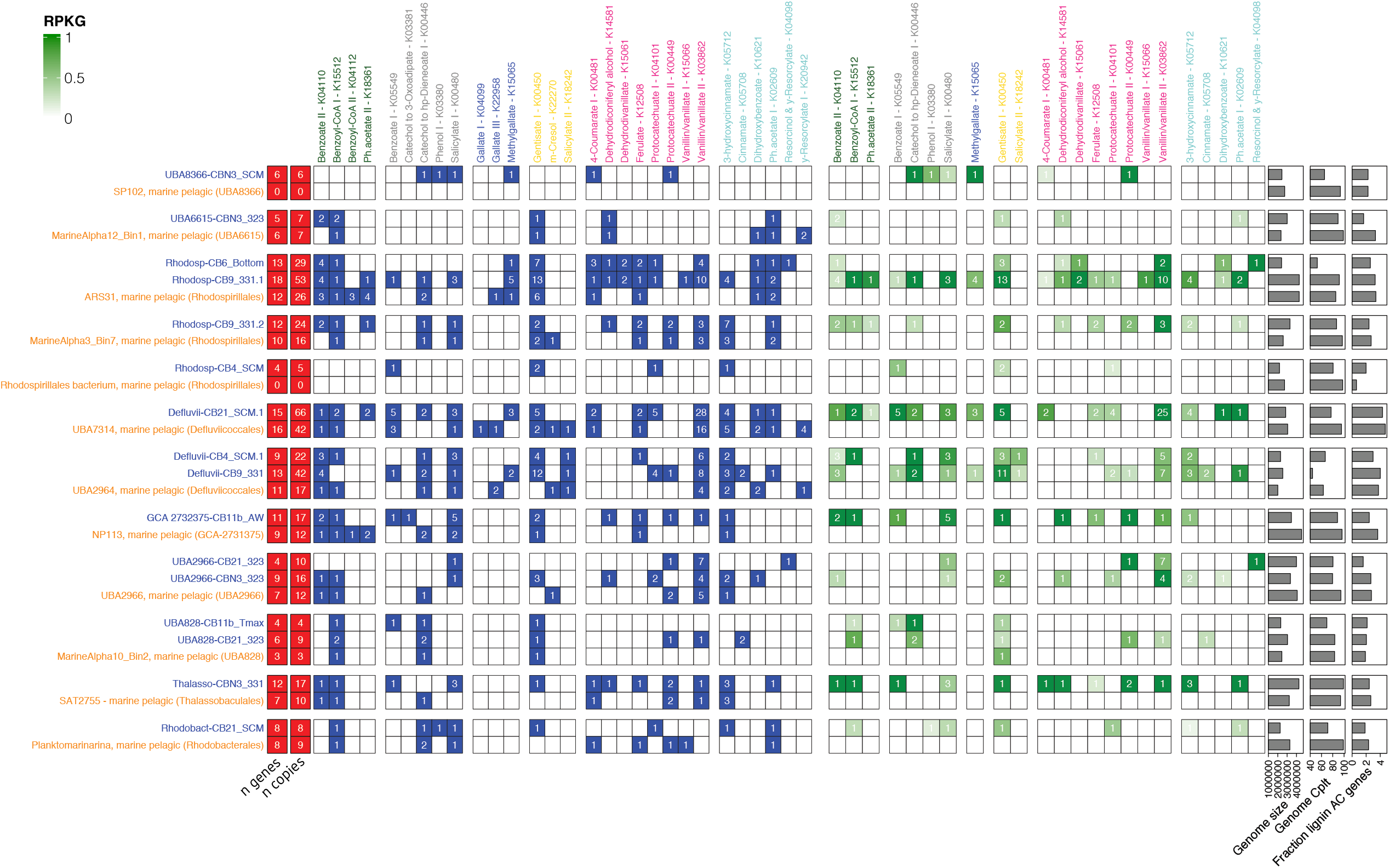
Comparative genomics of the the Arctic Ocean *Alphaproteobacteria* MAGs identified as most implicated in the degradation of aromatic compounds (blue) and their closest relative genomes (orange) from other studies. Left panel: graphical table of the presence (blue cells) and absence (white cells) of genes annotated with Kos marker of aromatic compound degradation pathways. Number in the cells represent the number of genes annotated with a KO. Right panel: graphical table of the mean expression (green cells) and absence of expression (white cells) of genes annotated with KOs marker of aromatic compound degradation pathways. Number in the cells represent the number of genes expressed.

Every single Arctic MAG contained at least one and up to seven marker genes that were not found in its closest related genome from other oceans (**Figure 7**). These included gentisate 1,2-dioxygenase (K00450) in Rhodosp-CB4_SCM, Defluvii-CB9_331, UBA2966-CBN3_323, Thalasso-CBN3_331 and Rhodobact-CB21_SCM, as well as vanillate O-demethylase (K03862) in Rhodosp-CB6_Bottom and Rhodosp-CB9_331.1 or benzoate CoA-ligase (K04110) in UBA6615-CBN3_323 and Rhodosp-CB9_331.2. These genes were the most abundant in the metagenomes and metatranscriptomes and most common throughout the Arctic Ocean microbiomes (**Figure 4a-c**). The three MAGs with the highest number of aromatic compound degradation gene copies (Defluvii-CB21_SCM.1, Rhodosp-CB9_331.1 and Defluvii-CB9_331) all possessed more copies of the vanillate O-demethylase (K03862) gene than their sister genomes. In addition, all of their sister genomes lacked the protocatechuate ring-opening genes (K04101 or K00449). A vast majority of the marker genes were also expressed in our Arctic Ocean MAGs (**Figure 7, right panel**), and we found only gentisate 1,2-dioxygenase (K00450) slightly expressed in the MarineAlpha10_Bin2 of the reference genomes.

## Discussion

### Aromatic compound degradation capacity of the Arctic Ocean microbiomes reflects the ability to degrade humic-rich DOM from terrestrial and sediment origin

The degradation of aromatic compounds emerged as a central metabolism of the Canada Basin microbiomes, which is consistent with a capacity to degrade the humic-rich DOM in the Arctic Ocean. We found that the aromatic compound degradation genes contributed more to the total metabolic capacity of the Arctic Ocean microbiome compared to other oceans, paralleling the higher concentrations of HS in the Arctic Ocean (Gonçalves-Araujo *et al*., 2016). This enhanced aromatic compound degradation capacity was associated with the humic-rich tOM layer of the Canada Basin (FDOMmax). Individually, most of the aromatic compound degradation genes were more abundant in the FDOMmax (**Figure 3a**), where humic-rich DOM concentrations are maximal, as evidenced by the distribution of the FDOM C1 fraction (**Figure 1c**). The lower abundance of aromatic compound degradation genes in the surface (**Figure 3a**) may be explained by a preference of the surface microbiomes to process the non-aromatic tOM fraction as sunlit waters are generally poor in aromatic compounds. This is supported by the higher contribution of the non-photoreactive (non-aromatic) FDOM C4 fraction and very low contribution of the aromatic C1 fraction in the surface (**Figure 1c**).

We found that the distribution and diversity of aromatic compound degradation genes in the Canada Basin matched the diversity of aromatic compounds expected from the Arctic Ocean watershed organic matter. Lignin is a polymer of three aromatic monolignols, which form the H, G and S unit (defined by 0, 1 and 2 methoxy group on the aromatic ring, respectively) when cross-linked. Coniferous trees dominate the boreal forest of the Arctic Ocean watershed, and are characterized by a high G/S ratio in their lignin polymer (Amon *et al*., 2012). Vanillin (G-unit) is therefore expected in the tOM of the riverine input to the Arctic Ocean. This has been shown in the Mackenzie River, the major river draining to the Canada Basin: the OM of suspended sediments in the Mackenzie was dominated by 1-methoxy aromatic compounds (G), among which vanillin and vanillate contributed the most (Goñi *et al*., 2000). The Mackenzie River also contained a significant contribution of benzoate derivatives (H) and smaller amounts of syringate derivatives (S). Our results showed that the genes involved in the degradation of vanillate, benzoate and methylgallate (syringate derivatives) were among the most abundant and expressed genes (**Figure 3a-b**) in the Canada Basin microbiomes, paralleling the aromatic compounds composition of the McKenzie River DOM input.

Sediments also contribute to the humic-rich OM reaching the Canada Basin, by exchanging OM with brine sinking along the shelf (Anderson and Macdonald, 2015). The OM on the Mackenzie shelf, bordering the Canada Basin, contained higher levels of vanillyl, syringyl and cinnamyl phenols than any other North American Arctic shelf (Goñi *et al*., 2013). The high abundance and expression of genes involved in the degradation of vanillate, syringate and 3-hydroxycinnamate (**Figure 3a-b**) suggest that the Canada Basin microbiomes access the variety of aromatic compounds from shelf sediments sources. This demonstrates that the Canada Basin microbiomes can access humic-rich DOM from terrestrial and sediment origin as growth substrates.

### *Rhodospirillales* are implicated as aromatic compound degraders in the Arctic Ocean

We showed that a few clades of *Alphaproteobacteria* were strongly implicated in aromatic compound degradation (**Figure 4, S8-S9**). Previous work in the Arctic Ocean showed that *Gammaproteobacteria* are associated with humic-rich DOM degradation. For example, in an experiment adding humic-rich OM derived from a thermokarst to coastal Arctic water (Chukchi sea) microbiomes, *Gammaproteobacteria* taxa *Colwelliaceae* (order *Alteromonadales*) rapidly dominated the microbial communities (Sipler *et al*., 2017). Similarily, *Alteromonadales* (genera *Glaciecola*, SAR92 clade) were associated with humic-rich riverine-derived OM consumption in an Arctic fjord (Lund Paulsen *et al*., 2019). These studies may seem incongruent with our results. However, these studies focused on the microbiomes of the surface waters only. In our study, we observed a significant *Gammaproteobacteria* signal in the taxonomy of aromatic compound degradation genes within the surface samples (**Figure 3, S7**), which supports earlier studies. In addition, the tOM used in previous experiments contained many other compounds than aromatic compounds, including more labile protein-like compounds. The *Gammaproteobacteria* in the surface could then be adapted to a fast consumption of pulses of labile OM, while the *Rhodospirillales* taxa of our study were more adapted to a slow degradation of more refractory and steadier amount of aromatic compounds-rich humic substances in the FDOMmax. This would be in line with the high variability in seasonal conditions and DOM concentrations in the surface waters of the Arctic Ocean compared to more stable DOM concentrations and conditions in the FDOMmax throughout the year (Davis and Benner, 2005; Tremblay *et al*., 2008).

All but one of the MAGs most implicated in aromatic compound degradation belonged to closely related *Alphaproteobacteria* clades, based on the GTDB. Based on NCBI taxonomy, all but one of these MAGs belonged to the *Rhodosprillales* order. Here we used the NCBI taxonomy to be able to relate our findings to previous reports in the literature. We concluded that the capacity to degrade aromatic compounds is phylogenetically concentrated in *Rhodosprillales* within the Arctic Ocean. In the global ocean, a few studies previously identified *Rhodospirillales* taxa in aromatic compound- and HS-degrading consortia. A *Rhodospirillales* strain (*Thalassospira profundimaris*) was one of six taxa isolated from the East China Sea surface microbiomes enriched with vanillic acid (Lu *et al*., 2020). This strain was able to grow on benzoic acid, 4-hydroxybenzoic acid, and to a lesser extent on syringate and ferulate. However, this *Rhodospirillales* strain is a member of a different clade (also including *Magnetospira* and *Magnetovibrio*) than the *Rhodospirillales* genomes we report in our study (**Figure 4a**). *Rhodospirillales* were also identified in flow-through experiments in which marine microbiomes were exposed to riverine HS as a sole carbon source (Rocker, Brinkhoff, *et al*., 2012). *Rhodospirillales* represented 6% of the taxa identified and were only reported in the low salinity experiment (14 PSU). Taxa were found in the *Thalassospira* (4 taxa) and *Thalassobaculales* (1 taxa) clades. We also reported *Thalassobaculales* genomes within the taxa implicated in aromatic compound degradation. However, the genomes we reported were located in the water column at salinities > 30 PSU. The studies reporting *Rhodospirillales* focused only on the surface water microbiomes, while we investigated the whole water column. The focus on surface waters in other studies is usually based on the assumption that aromatic compounds and HS have a terrestrial origin and are transported to the ocean with freshwater input, therefore concentrated in the surface layers. The focus on surface waters microbiomes may therefore explain why *Rhodospirillales* have not yet been reported as most implicated in aromatic compounds and HS degradation within the ocean. Based on the distribution of HS in the Arctic Ocean, we investigated the whole water column and specifically the FDOMmax, which allowed us to identify *Rhodospirillales* as strongly implicated in aromatic compounds degradation. Further work within other oceans, as well as experimental work using HS as sole carbon sources in the microbiomes of the FDOMmax will be necessary to fully elucidate the role of *Rhodospirillales* in the degradation of HS in the Arctic Ocean, and the global ocean.

### Evolutionary adaptation of *Rhodospirillales* in the Arctic Ocean

The phylogenetic divergence of our *Rhodospirillales* MAGs from relatives in other oceanic regions and their restricted distribution to the Arctic Ocean suggests the Arctic populations are evolutionarily adapted to life in the Arctic Ocean. The high number of aromatic compound degradation gene copies in the MAGs compared to their closest relatives in other oceans suggest that the capacity to use aromatic compounds as a growth substrate played a role in their evolutionary adaptation. The disproportionately high amount of HS in the FDOMmax may then act as a selective pressure on these MAGs. Previous studies have demonstrated evolutionary adaptation of microbial MAGs restricted to the Canada Basin: a new *Methylophilaceae* clade (Ramachandran, McLatchie and Walsh, 2021) as well as SAR11 (Kraemer *et al*., 2020) and *Chloroflexi* ecotypes (Colatriano *et al*., 2018). The *Methylophilaceae* clade evolved via a freshwater to marine transition, highlighting the importance of the terrestrial-marine interface in shaping the Canada Basin microbiomes. However, HS does not appear as the main selective pressure for *Methylophilaceae* as their distribution is restricted to the surface. The Arctic SAR11 and *Chloroflexi* clades were restricted to the SCM and FDOMmax, where humic-rich DOM is enriched within the Canada Basin. The *Chloroflexi* Arctic ecotype was replete with aromatic compound degradation genes, some of these acquired by lateral gene transfer from terrestrial taxa. This *Chloroflexi* ecotype was found within the water masses rich in HS (FDOMmax), similarly to the *Rhodospirillales* MAGs of our study. The preference for humic-rich water masses coupled to an enhanced capacity to degrade aromatic compounds in *Chloroflexi* and *Rhodospirillales* suggest that the ability to use aromatic compounds as growth substrate provides an evolutionary advantage in the humic-rich environment of the Canada Basin FDOMmax.

## Conclusion

The dissolved organic matter of the Arctic Ocean is characterized by a disproportionately high contribution of HS compared to other oceans. With the increasing terrestrial input of humic-rich OM to the Arctic Ocean as a result of escalating permafrost thawing and river runoff, it is predicted that the contribution of HS to the Arctic Ocean OM will increase (Gueguen and DeFrancesco, 2021). The fate of this carbon is important to consider with respect to changing biogeochemical cycles of the Arctic Ocean. In this study, we showed that the metabolic pathways involved in the degradation of HS were widespread, abundant and expressed in the microbiomes of the Canada Basin. The capacity to degrade humic-rich OM in the Arctic Ocean microbiomes was enhanced compared to the microbiomes of the global ocean in the upper water column. The diversity and distribution of the aromatic compound degradation machinery revealed that the Arctic Ocean microbiomes were equipped to use OM from terrestrial sources as growth substrates. We identified that the aromatic compound degradation capacity was concentrated phylogenetically in *Rhodospirillales*. The phylogeny, comparative genomics and biogeographic distribution of these *Rhodospirillales* suggest an evolutionary adaptation driven by the disproportionately high amount of HS in the Arctic Ocean. Overall, this study demonstrates that the Arctic Ocean microbiomes are capable of processing OM of terrestrial origin. Our study predicts that OM of terrestrial origin can be remineralized in the Arctic Ocean and that *Rhodospirillales* will gain importance as tOM inputs continue to increase in the Arctic Ocean.

## Methods

### Sampling, DNA and RNA extraction

Samples were collected in September 2017 during the Joint Ocean Ice Study cruise to the Canada Basin. We analyzed 22 metagenomes and 25 metatranscriptomes generated from samples collected across the water column of the Canada Basin. Eight specific water masses were sampled: the surface mixed layer (surface: 5 m and 20 m depth) characterized by fresher water due to riverine input and ice melt, the subsurface chlorophyll maximum (SCM), in the halocline (FDOMmax at salinity of 32.3 and 33.1 PSU, referred as 32.3 and 33.1), and deeper water from Atlantic origin at the temperature maximum (referred as Tmax), 1000 m depth (Atlantic water, further referred as AW) and 10 or 100 m above the bottom (further referred as bottom).

We filtered 14L of seawater for DNA samples and 7L of seawater for RNA samples sequentially through a 3 μm pore size polycarbonate track etch membrane filter (AMD manufacturing, ON, Canada) and a 0.22 μm pore size Sterivex filter (Millipore, MA, USA). Filters were stored in RNALater (ThermoFischer, MA, USA), and kept frozen at -80^0^C until processing in the lab. DNA was extracted following the method described in (Colatriano and Walsh, 2015). Briefly, the preservation solution was expelled and replaced by a SDS solution (0.1 M Tris-HCl pH 7.5, 5% glycerol, 10 mM EDTA, 1% sodium dodecyl sulfate) and incubated at room temperature for 10 min and then at 95^0^C for 15 min. The cell lysate was then centrifuged at 3,270 x g. Proteins were removed by precipitation with MCP solution (Lucigen, WI, USA) and the supernatant was collected after centrifugation at 17,000 x g for 10 min at 4^0^C. DNA was precipitated with 0.95 volume of isopropanol and rinsed twice with 750 μL ethanol before being air dried. The DNA was resuspended in 25 μL of low TE buffer, pH 8 (10 mM Tris-HCl, 0.1mM EDTA) and stored at -80^0^C.

The RNA extraction procedure was adapted from the mirVana RNA extraction kit (ThermoFisher, MA, USA). RNAlater was expelled from the Sterivex and replaced by 1.5 mL of Lysis buffer and Sterivex was vortexed. 150 μL of miRNA homogenate were added, the Sterivex vortexed and incubated on ice for 10 min. The cell lysate was expelled from the Sterivex, 0.9x the volume of acid-phenol-chloroform was added and the solution was vortexed for 30-60 sec. The mix was centrifuged at 10,000 x g for 5 min, and the top aqueous phase gently removed and transferred to a fresh tube. 1.25 volume of 100% ethanol was added to the aqueous phase and vortexed to mix. The mix was filtered through mirVana Filter Cartridges by centrifugating at 10,000 x g for 10 s, and the flow through discarded. The RNA was rinsed with 700 μL of Wash Solution 1 and then with 500 μL Wash solution 2/3 by centrifugating at 10,000 x g for 10 sec. RNA was then eluted with 50 μL of Elution solution (0.1 M EDTA) warmed at 95^0^C. 700 μL of RTL buffer and 500 μL 100% ethanol were added to the RNA suspension and the suspension was centrifuged for 15 sec at 10,000 x g on a RNeasy MinElute column. RNA was washed first with RPE buffer by centrifuging 500 μL for 15 sec at 10,000 x g and then 80% ethanol for 2 min at 10,000 x g. The empty column was then centrifuged at 12,000 x g for 5 min to discard the excess liquid. The RNA was finally eluted by centrifugation of 28 μL and then 10 μL of RNase free water for 1 min at 12,000 x g and stored at -80^0^C.

### Dissolved organic matter samples collection, analysis of fluorescence measurements

DOM samples were collected in Niskin bottles mounted on a conductivity-temperature-depth rosette profiler and immediately filtered using pre-combusted 0.3um glass fiber filters (GF75, Advantec) into pre-combusted amber glass vials. Fluorescence spectra were measured using a Fluoromax 4 Jobin Yvon fluorometer (DeFrancesco and Guéguen, 2021). Parallel factor analysis was applied to decompose the fluorescence signal into their main components following the procedures outlined in Murphy *et al* (Murphy *et al*., 2013). The PARAFAC model validated 7 components including 5 humic-like (C1-C2, C4-C5, and C7) and 2 protein-like components (C3 and C6) in 4,483 samples collected from surface to 10 m above bottom sediment in the Canada Basin between 2007 and 2017 (Gueguen and DeFrancesco, 2021).

### Metagenomic sequencing, assembly and annotation

Sequencing, assembly and annotation were performed by the Joint Genome Institute (CA, USA). Each individual metagenome and metatranscriptome were sequenced on the Illumina NovaSeq platform, generating paired-end reads of 2×150 bp for all libraries. Single assemblies were created for each individual sample using SPAdes (Bankevich *et al*., 2012) with kmer sizes of 33,55,77,99,127 bp. Gene prediction and annotation was performed using the DOE Joint Genome Institute Integrated Microbial Genomes Annotation Pipeline v.4.16.5 (Huntemann *et al*., 2016).

### Building of EC and KO abundance matrices

The number of copies of genes annotated to each Enzyme Commission (EC) number or KEGG Orthology (KO) number in a metagenome or metatranscriptome was calculated by summing the depth of coverage of all genes or transcripts annotated with this EC or KO. To obtain the final EC and KO abundances (number of gene or transcript copies/10^6^ reads for this EC or KO) the total number of copies were then normalized by the library size (number of reads) with the TMM method (Robinson and Oshlack, 2010), using the calcNormFactors function of the edgeR package in *R* (Robinson, McCarthy and Smyth, 2009). Genes were assigned to a total of 3,102 EC numbers and 12,018 KO identifiers for the metagenomes and 2,830 EC numbers and 10,556 KO identifiers for the metatranscriptomes. Before multivariate analysis, EC and KO abundances were transformed with a Hellinger transformation (decostand function from the R vegan package (J *et al*., 2007)).

### Multivariate analysis

Non-metric multidimensional scaling (NMDS) was performed on the EC number abundance matrices with the metaMDS function from the *R vegan* package, using two dimensions and the Bray-Curtis dissimilarity metric. Non-negative matrix factorization (NMF) was performed with the nmf function from the NMF package in R (Gaujoux and Seoighe, 2010). NMF decomposes the abundance matrix into two matrices: a coefficient matrix that describes the overall structure of the abundance matrix with a limited number of descriptors (called sub-metagenomes and sub-metatranscriptomes in this study, their number being the rank), and a basis matrix the provides the weights of each original descriptors (EC number) on the new descriptors (sub-metagenomes, sub-metatranscriptomes). The advantage of NMF is to directly link the overall structure of the abundance matrix to the individual elements (EC number) driving this structure. We first performed the NMF analysis with rank values ranging from 3 to 7, 100 runs, and various algorithms (“nsnmf “, “Brunet”, “KL”). We obtained the optimal results for the nsNMF algorithm, random seed of the factorized matrices, and a rank value of 4. We performed the final analysis with 200 runs, rank of 4, random seed and nsNMF algorithm.

### Calculation of gene and pathway indices

The indices were calculated for EC number annotated from metagenomes and metatranscriptomes by combining two methods described in Jiang *et al*. (Jiang *et al*., 2012) and Kim *et al*. (Kim and Park, 2007). We first used the EC number abundance matrices (annotated from metagenomes and metatranscriptomes) and the coefficient matrices (SMG/SMT x samples) to calculate both the spearman correlation coefficient and the multidimensional projection between all pairs of EC number annotated from metagenomes (EC_MG_) and SMG as well as all pairs of EC number annotated from metatranscriptomes (EC_MT_) and SMT. The spearman correlation coefficient between an EC_MG_/EC_MT_ (*i*) and a SMG/SMT (*k*) _*i,k*_ was calculated using the abundance profile of a EC_MG_/EC_MT_ and a SMG/SMT along all the samples. The multidimensional projection between an EC_MG_/EC_MT_ and a SMG/SMT was calculated as the cosine of the angle between the vectors represented by an EC_MG_/EC_MT_ abundance in the samples space and the vector represented by SMG/SMT in the samples space. The abundance profiles of EC_MG_/EC_MT_ and SMG/SMT were first normalized, and the multidimensional projection was calculated as:

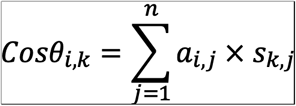

Where *Cosθ*_*i,k*_ is the multidimensional projection between the EC_MG_/EC_MT_ *i* and the SMG/SMT *k, n* is the number of samples, *a*_*i,j*_ is the normalized abundance of the EC_MG_/EC_MT_ *i* in the sample *j*, and *s*_*k,j*_ is the normalized abundance of the SMG/SMT *k* in the sample *j*. We then used the basis matrix to calculate the score of each EC_MG_/EC_MT_:

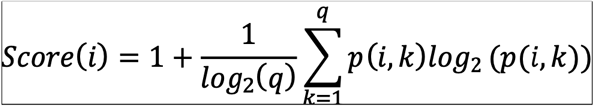

Where *i* is the EC_MG_/EC_MT_, *q* is the number of SMG/SMT (4 in our study), *k* is the SMG/SMT, *p(i,k)* is the probability of finding the EC_MG_/EC_MT_ *i* in the SMG/SMT *k*

We calculated the final EC index (annotated from metagenome and metratranscriptome) on each SMG/SMT by multiplying the spearman correlation coefficient, cos theta and the EC score:

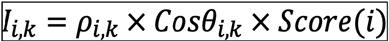

This allowed us to calculate an index for each pair of EC_MG_/EC_MT_ and SMG/SMT.

### Aromatic compound degradation gene and pathway selection

To select the metabolic pathways and enzymes involved in the degradation of lignin-derived aromatic compounds, we first surveyed the literature for the various lignin breakdown compounds reported to be degraded by bacteria (Bugg *et al*., 2011; Kamimura *et al*., 2017; Brink *et al*., 2019). We then retrieved the MetaCyc (https://metacyc.org) pathways that were involved in the degradation of these compounds, as well as the EC numbers involved in these pathways. For analysis of marker genes for each pathway, we first selected one key reaction in each pathway. We then retrieved genes annotated with KO numbers corresponding to the EC numbers associated with the key reactions. We first chose reactions involved in aromatic ring-opening, aromatic ring-oxidation, and aromatic ring-reduction steps. If the EC numbers of these reactions were not specific to a pathway, we chose reactions involved in the addition of CoA to the aromatic ring or reactions involved in oxidoreduction steps of the side chains of the aromatic ring. If 2 aromatic compounds degradation pathways possessed the same EC number associated with the selected key reaction, and if this EC number was not found on any other Metacyc pathways, we used the EC as marker for only one of the two pathways. We could not retrieve any marker EC number for several pathways.

### Calculation of the fraction of genes involved in the degradation of aromatic compounds within the pool of metabolic genes

To calculate the percentage of the gene pool associated with the degradation of aromatic compounds in each sample, we summed the total number of genes copies annotated with EC number within our selection of aromatic compound degradation pathways. We then divided this number by the total number of gene copies annotated with EC numbers.

### Statistical analyses

To compare the means of EC indices distribution as well as the percentage of aromatic compounds degradation genes in metagenomes, we first performed a permutational ANOVA (PERMANOVA) using the perm.anova function of the RVAideMemoire package in R. When the p-value of the PERMANOVA test was less than 0.05, we performed pairwise comparison between groups with Student t-test, using the PERMANOVA residuals variance as the variance for the Student t-tests. Two groups were considered different if their p-value was less than 0.05.

### Calculation of aromatic compound degradation pathways completeness

Pathway completeness in a sample was calculated by dividing the number of EC numbers belonging to this specific pathway and present in a sample by the total number of EC number of this specific pathway. The marker genes abundance was obtained using the normalized abundance of a specific KO as calculated to obtain the KO abundance matrices (see above).

### Estimation of the fraction of taxa harboring aromatic compound degradation genes

The estimated percentage of genomes harboring marker genes in a sample was calculated by dividing the total number of gene variants annotated with the KO of interest by the total number of gene variants annotated with the KO corresponding to the *recA* gene (K03553).

### Taxonomic assignment of aromatic compound degradation genes

Genes annotated with KO identified as marker for selected lignin-derived aromatic compound degradation pathways were grouped by water column feature and dereplicated with CD-hit (v.4.6) (Li and Godzik, 2006) at 95% identity. The dereplicated set of genes were searched against the NCBI nr protein database (downloaded 21/08/27) using DIAMOND (v.0.9.30.131) (Bağcı, Patz and Huson, 2021). To assign a taxonomic identity to these genes, the DIAMOND (v.0.9.30.131) output was imported in MEGAN (Bağcı, Patz and Huson, 2021) using the January 2021 mapping file (“megan-map-Jan2021.db”). The lowest common ancestor parameters were set at minimum e-value of 1×10^−20^ and at top percent of 1%. The file containing taxonomic identity of the genes was then exported from MEGAN and processed with a custom-made *R* script.

### Metagenome binning

Metagenomic binning was performed on each individual assembly with Metabat2 (v.2.12.1) (Kang *et al*., 2019) using scaffold longer than 2500 bp. Contamination and completeness of the metagenome-assembled genomes (MAGs) were estimated with CheckM (v.1.0.7) (Imelfort *et al*., 2015). MAGs greater than 30% completeness and less than 10% contamination were selected for further analysis. Phylogenetic placement of MAGs was performed based on the concatenation of 120 conserved genes for bacteria and 122 conserved gene for archaea using the Genome Database Taxonomy Database toolkit (GTDB-Tk – v.1.3.0) (Parks *et al*., 2018; Chaumeil *et al*., 2020).

### Metabolic reconstruction and MAG selection based on the capacity to degrade aromatic compounds

To select MAGs enriched in aromatic compound degradation capacity, we selected all the genes annotated with EC numbers belonging to pathways involved in the degradation of aromatic compounds within the MAGs. As ring-fission pathways can be involved in the degradation of non-lignin aromatic compounds, we used only the funneling pathways to select for MAGs enriched in the degradation of lignin aromatic compounds. For each MAG, we calculated the completeness of all funneling aromatic compound degradation pathways. We considered a pathway complete if a MAG contained genes annotated with all the EC number of this pathway. The pathway completeness percentage was obtained by dividing the number of EC numbers involved in a pathway within a MAG by the number of reactions of this pathway. This number was normalized by the MAG completeness. For each MAG, we then calculated the median of all pathway completeness. Based on the distribution of the medians of all MAGs, we selected 4% as the median threshold above which a MAG was selected as having a high capacity to degrade aromatic compounds. We obtained a total of 46 MAGs. We calculated the average nucleotide identity (ANI) between these 46 MAGs using fastANI (v.1.3) (Jain *et al*., 2018) and grouped the MAGs with an ANI>95% as the same genomospecies, obtaining a total of 22 genomospecies.

### Phylogenetic analyses of MAGs

To reconstruct the phylogeny of the 38 *Alphaproteobacteria* MAGs (16 genomospecies), we first manually investigated their phylogenetic placement within the GTDB. For each of our selected MAGs, we picked the most closely related genomes from the GTDB, as well as genomes representative of distinct families. We then reconstructed a phylogeny with our selected MAGs and the selected genomes from GTDB using concatenation of 120 conserved genes and FastTree (Chaumeil *et al*., 2020)

### Metagenome and metatranscriptome fragment recruitment

In order to evaluate the abundance and overall expression of our selected MAGs across the samples, we mapped the reads of the metagenomes and metatranscriptomes to our selected MAGs using bbmap (v.35) and a minimum sequence identity of 98%. We then calculated final RPKG values (reads per MAG kilo base pairs per metagenome giga base pairs), by dividing the total number of reads mapped to each MAG, by the size of the MAG (kbp) and the size of the metagenome/metatranscriptome (Gbp).

In order to evaluate the distribution of selected *Alphaprotebacteria* MAGs and their most closely related reference genomes across oceans, we performed fragment recruitment as in (Kraemer *et al*., 2019). The 16 MAGs representative of the genomospecies enriched with the capacity to degrade aromatic lignin moieties, as well as 12 closely related reference genomes were searched against the metagenomic dataset using blastall (v.2.2.25) (e-value=0.00001). The recruited reads were extracted from the metagenomes and searched against a database consisting of the concatenation of all 28 genomes (16 Arctic *Alphaproteobacteria* MAGs and 12 reference genomes) using blastall (v.2.2.25). We selected the best hit, filtered for a minimum of 100 bp alignment and 98% sequence identity. We then calculated the RPKG values by normalizing the number of reads recruited by kilobase of genome and gigabase of metagenome.

### Annotation of publicly available genomes

Gene sequences were retrieved from publicly available genomes at NCBI (**Table S4**) and translated to proteins. Ribosomal RNA genes were predicted in Infernal v. 1.1.2 (Nawrocki & Eddy, 2013) against Rfam v. 14.2 (Kalvari *et al*., 2021). Gene functions were annotated in KofamScan using default settings and a bitscore-to-threshold ratio of 0.7 (Aramaki *et al*., 2020).

## Supporting information

Tabls S6

## Availability of data and materials

The metagenomic data generated in this study are available in the Integrated Microbial Genomes database at the Joint Genome Institute at https://img.jgi.doe.gov, GOLD Project ID: Gs0134626. Metagenome-assembled genome projects will be deposited at DDBJ/ENA/GenBank under the Bioproject XXXXXX and accession numbers XXXXXX. For the purpose of review, the MAG datasets (.fa, .faa, and .gff files) supporting the conclusions of this article are included as supporting documents.

## Authors’ contributions

D.A.W designed the study. T.G. generated the metagenomic and metatranscriptomc data and performed the bioinformatic analyses. T.G. and D.A.W. wrote the manuscript. V.O. performed the functional annotation of the genomes. C.G. collected the samples and contributed to the analysis of the FT-ICRMS data. All authors reviewed the manuscript.

## Acknowledgments

The data were collected aboard the CCGS Louis S. St-Laurent in collaboration with researchers from Fisheries and Oceans Canada at the Institute of Ocean Sciences and Woods Hole Oceanographic Institution’s Beaufort Gyre Exploration Program and are available at http://www.whoi.edu/beaufortgyre. We would like to thank both the Captain and crew of the CCGS Louis S. St-Laurent and the scientific teams aboard.

