## Supplementary figures and images for "Degradation pathways for organic matter of terrestrial origin are widespread and expressed in Arctic Ocean microbiomes"

### Figure S1

#### Figure S1

##### Metagenome

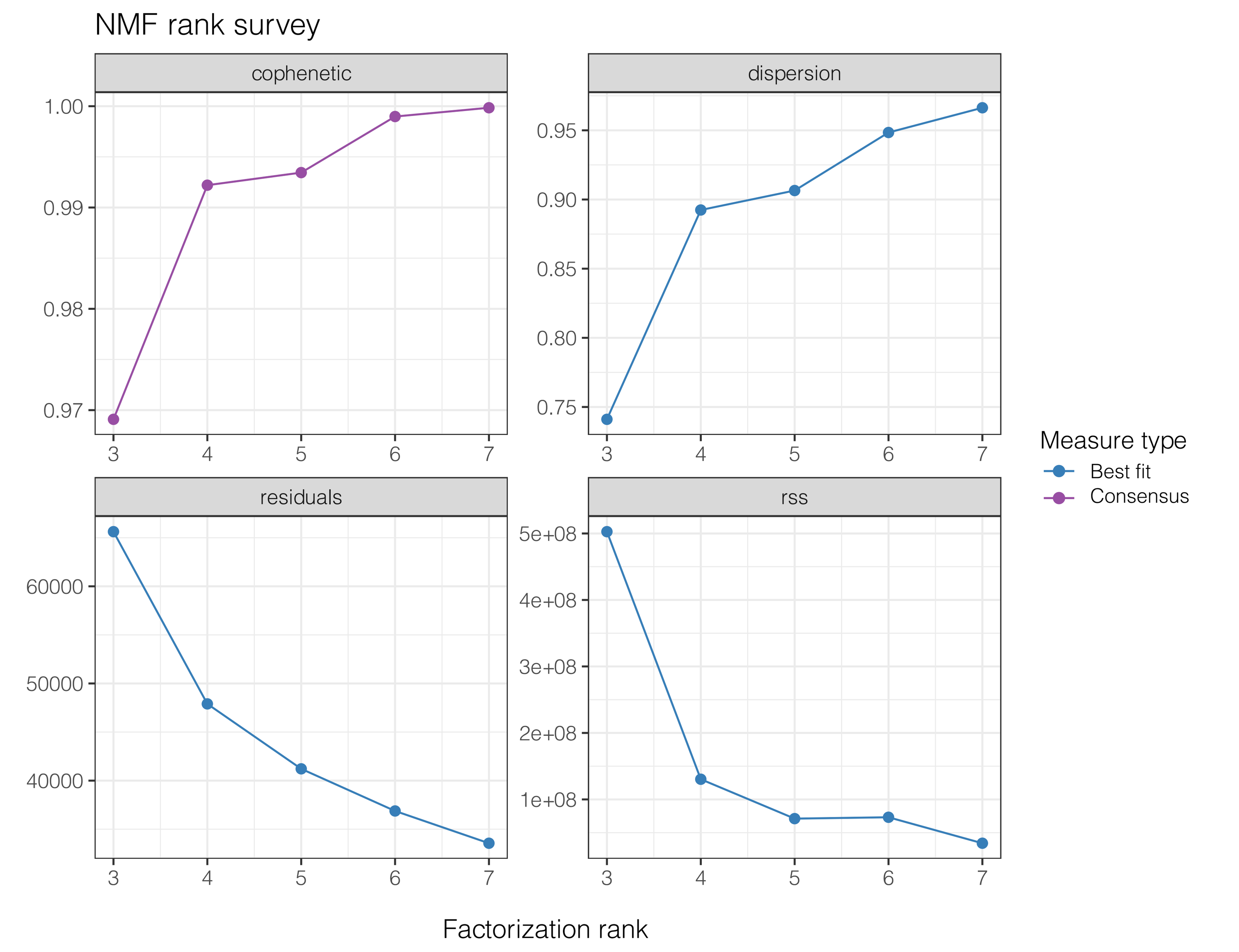


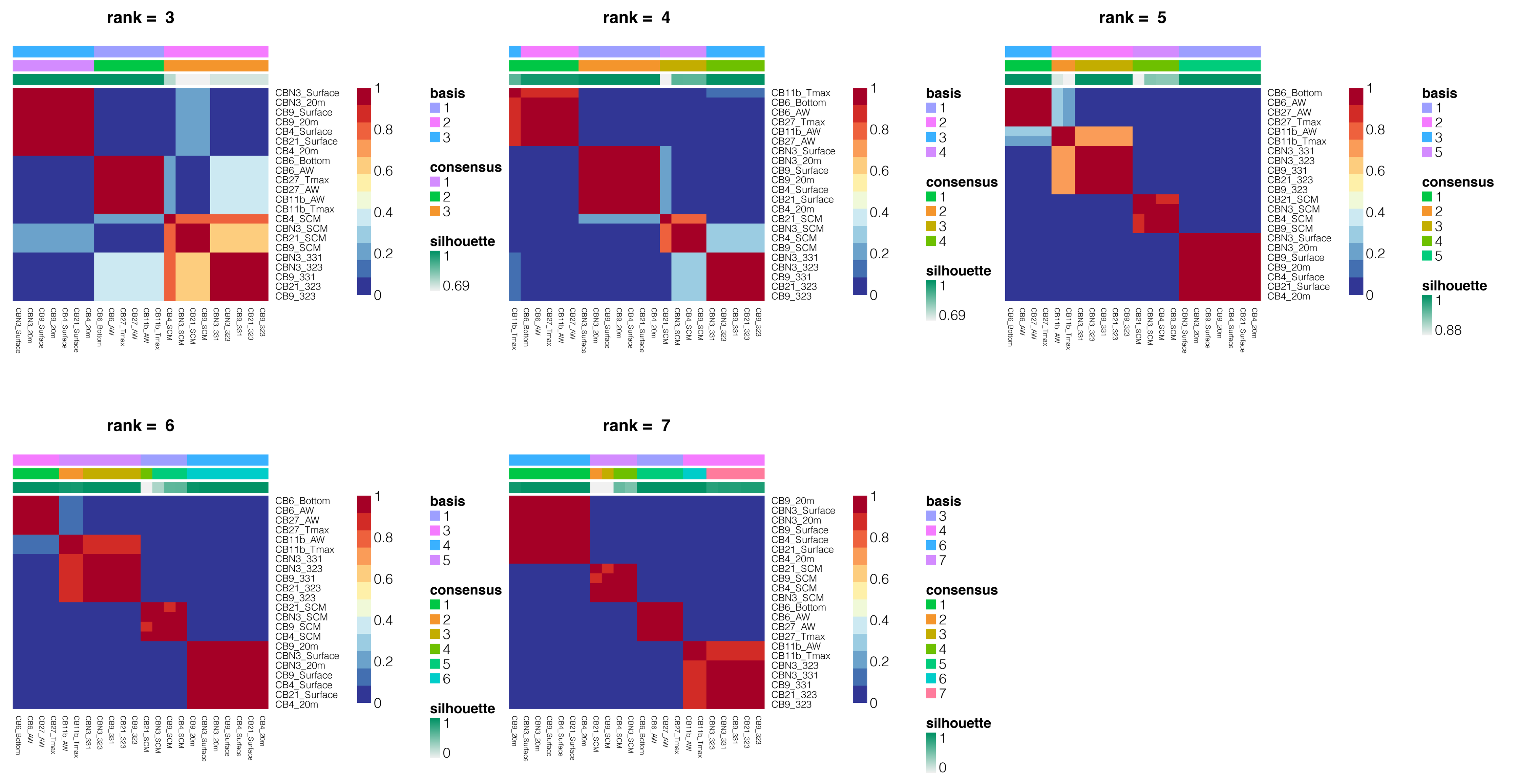


##### Metatranscriptome

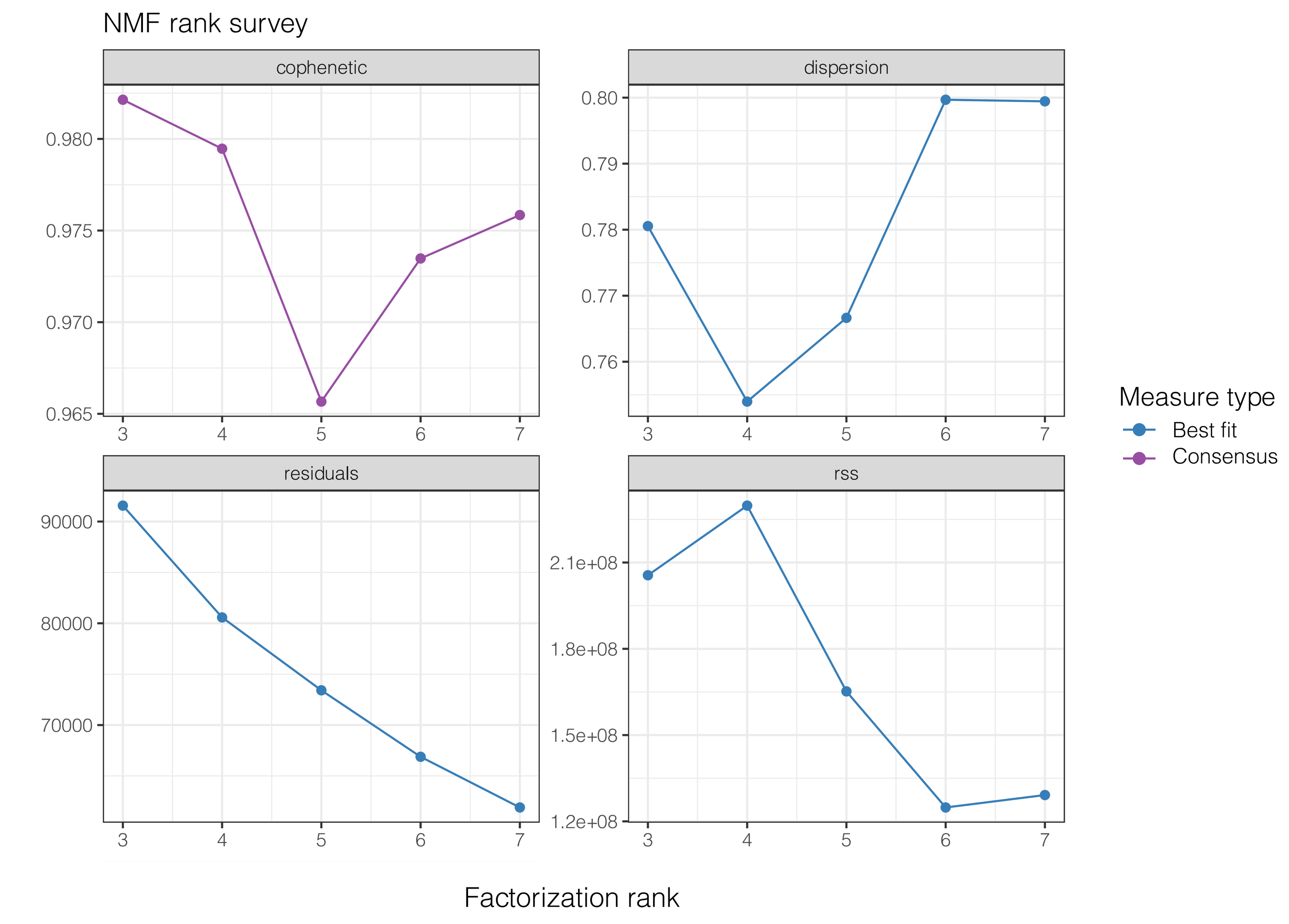


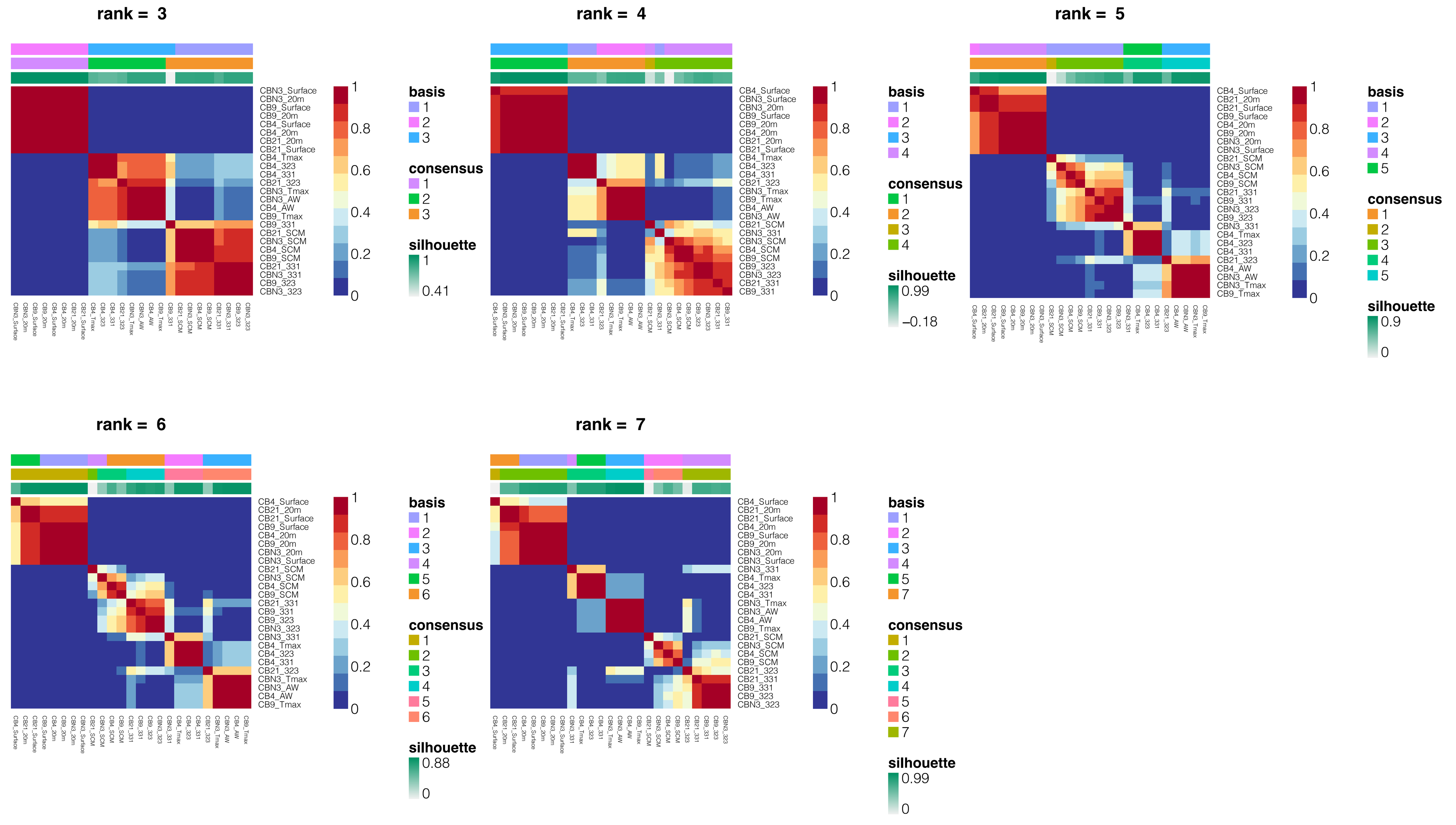

### Figure S2

#### Figure S2

##### Basis matrix

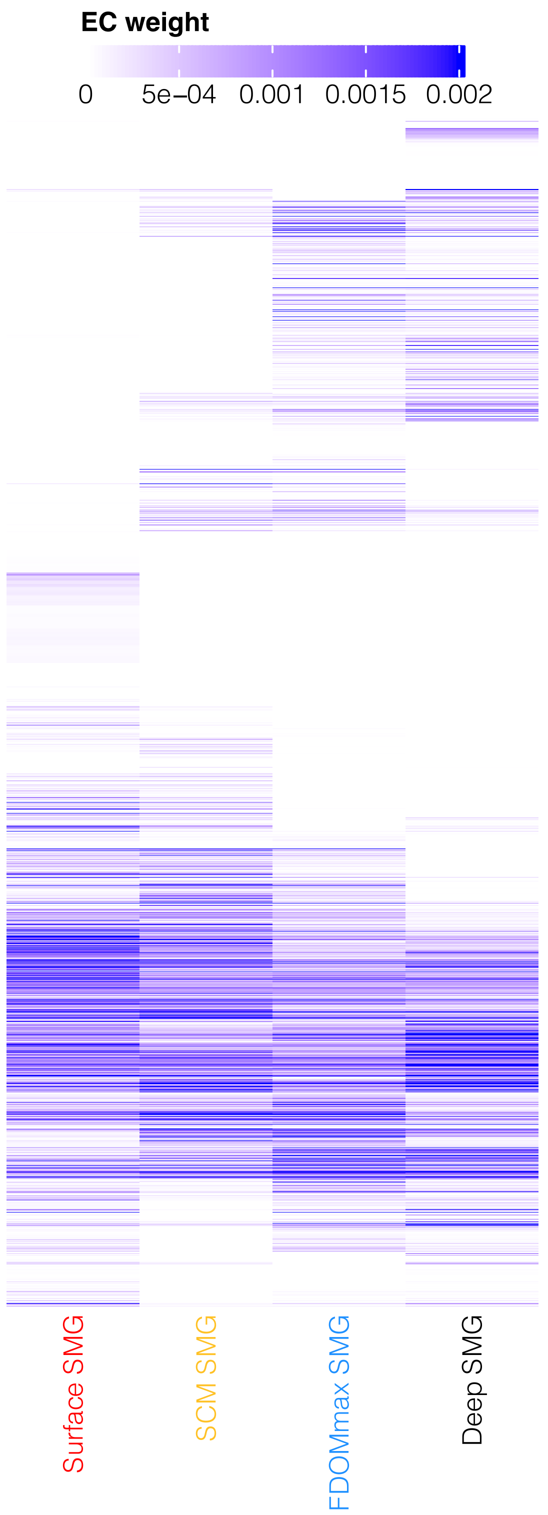


##### Coefficient matrix

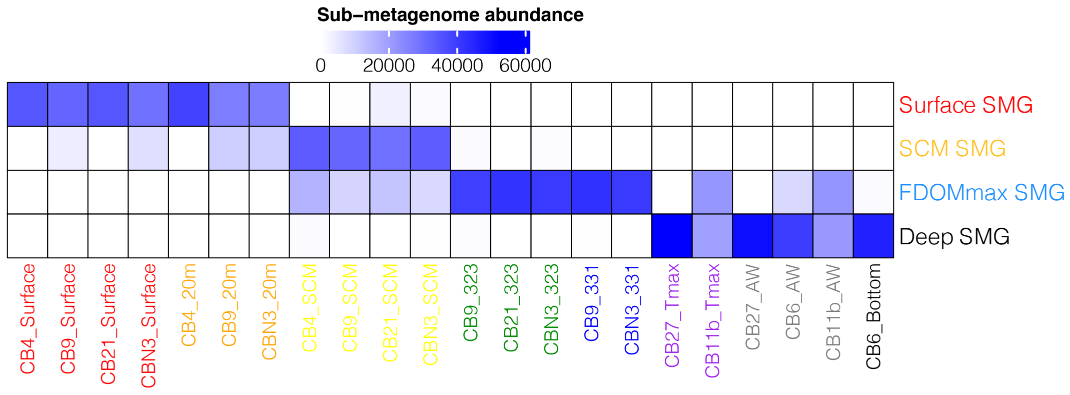

### Figure S3

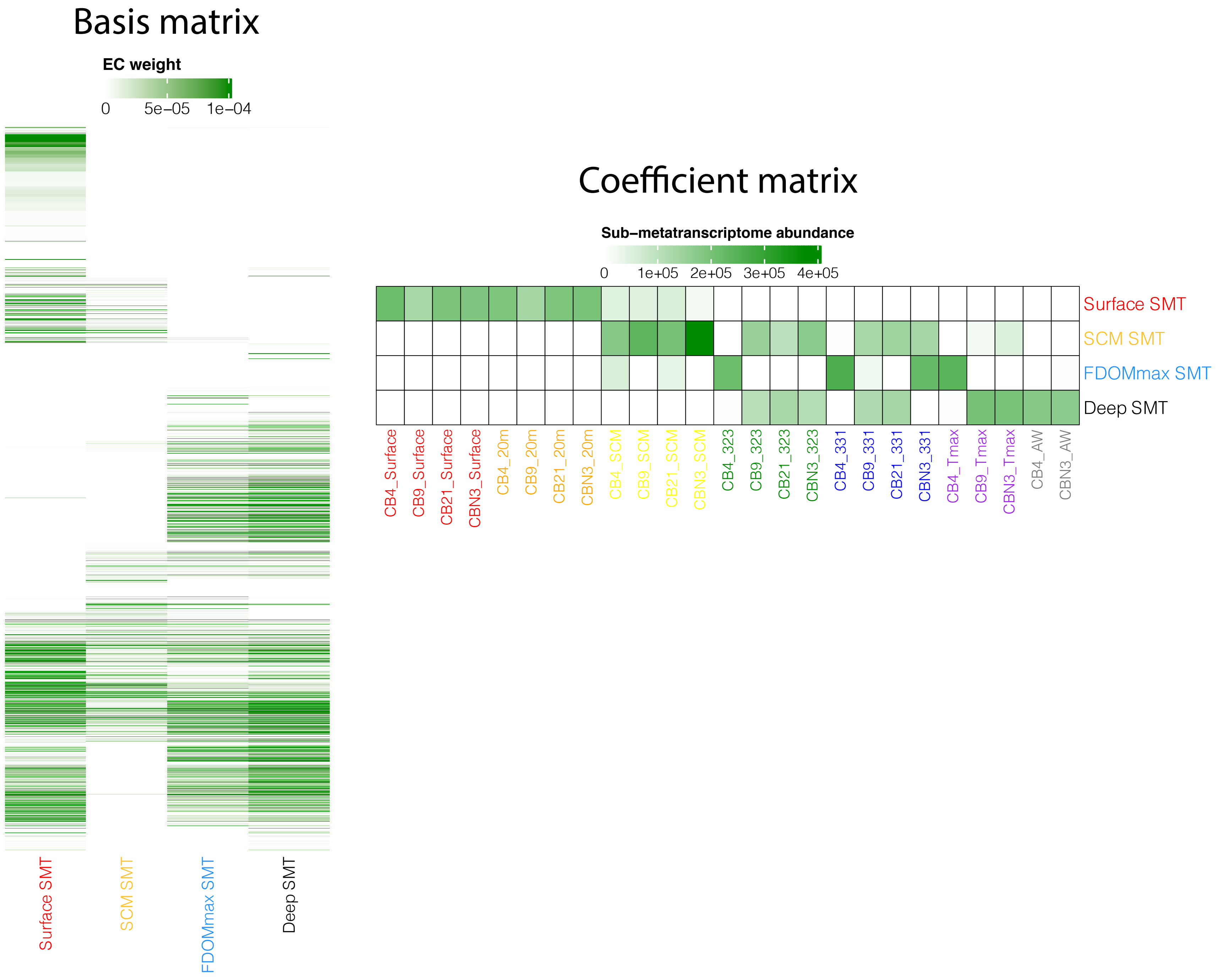

### Figure S4

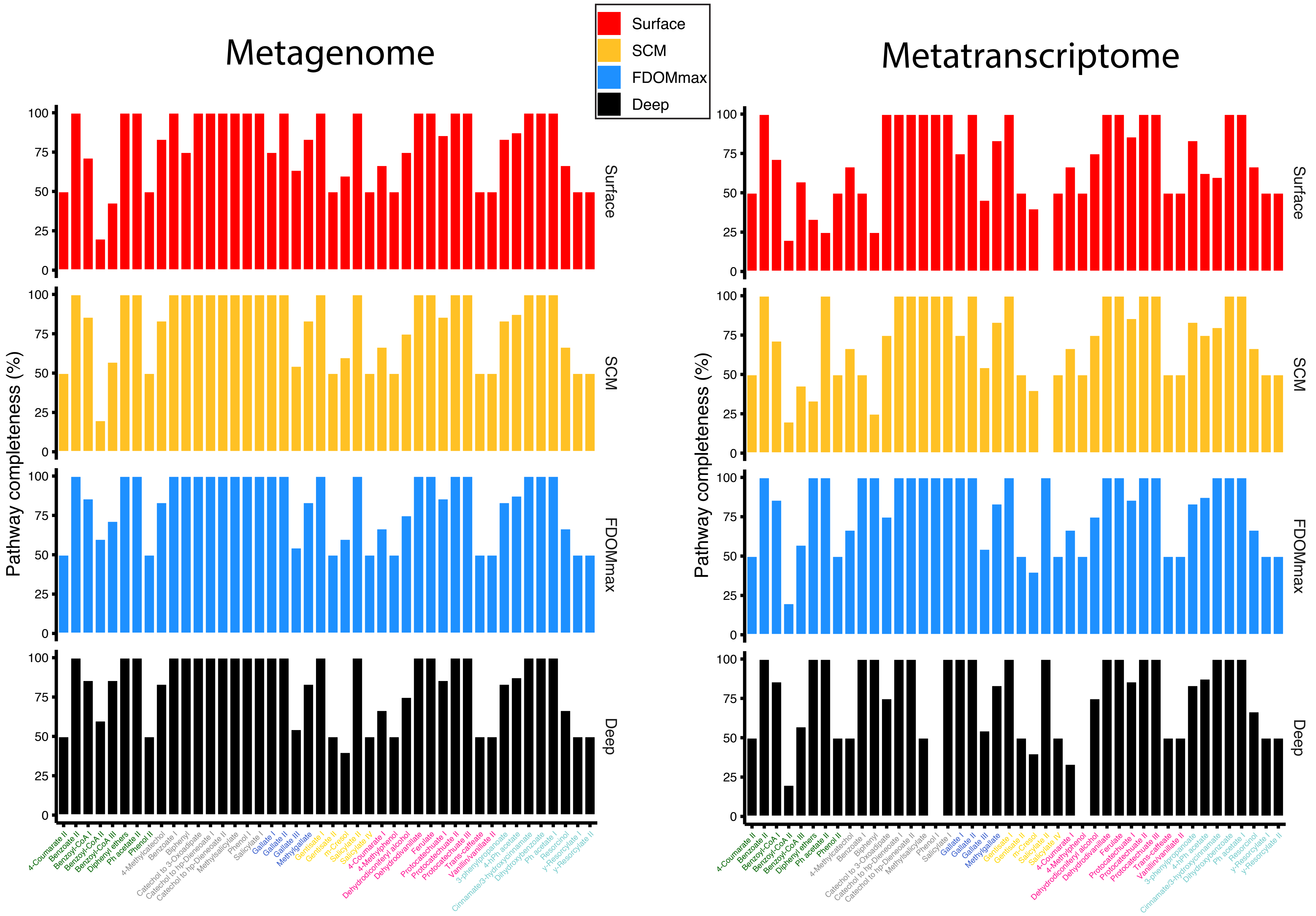

### Figure S5

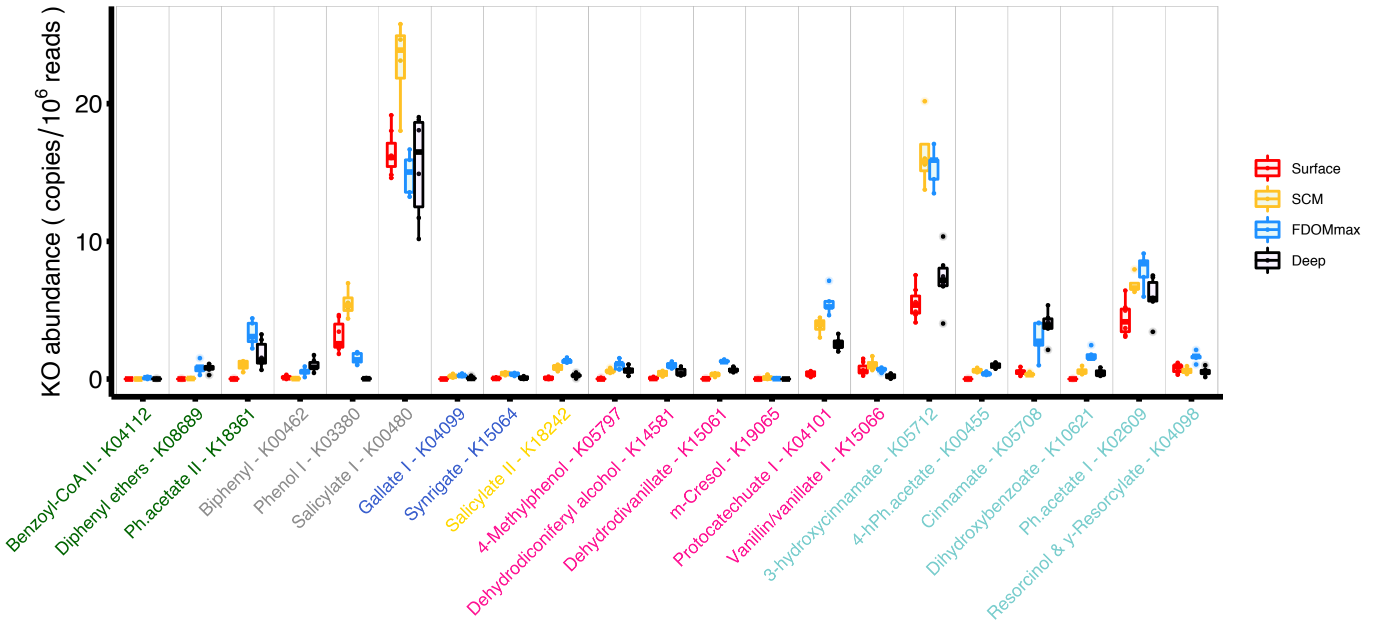

### Figure S6

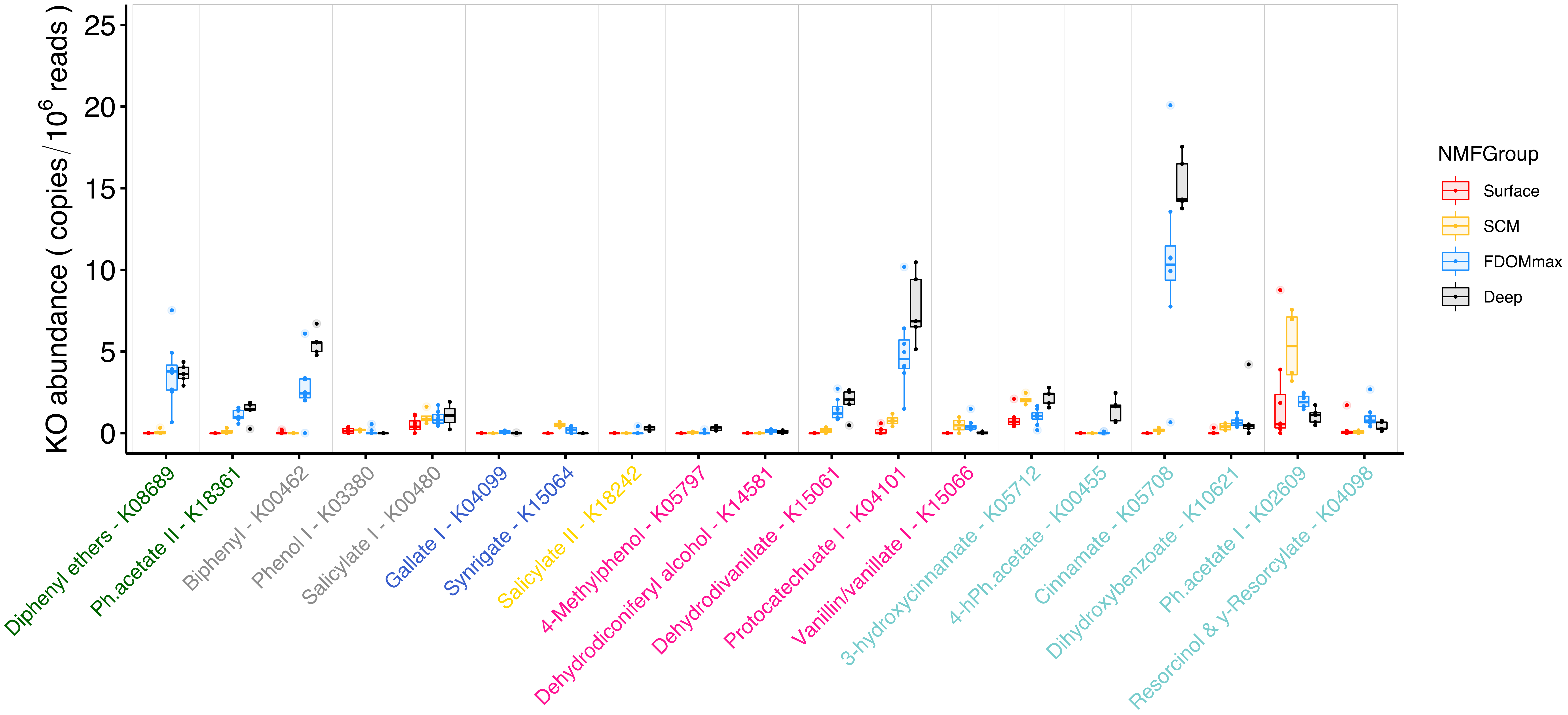

### Figure S7

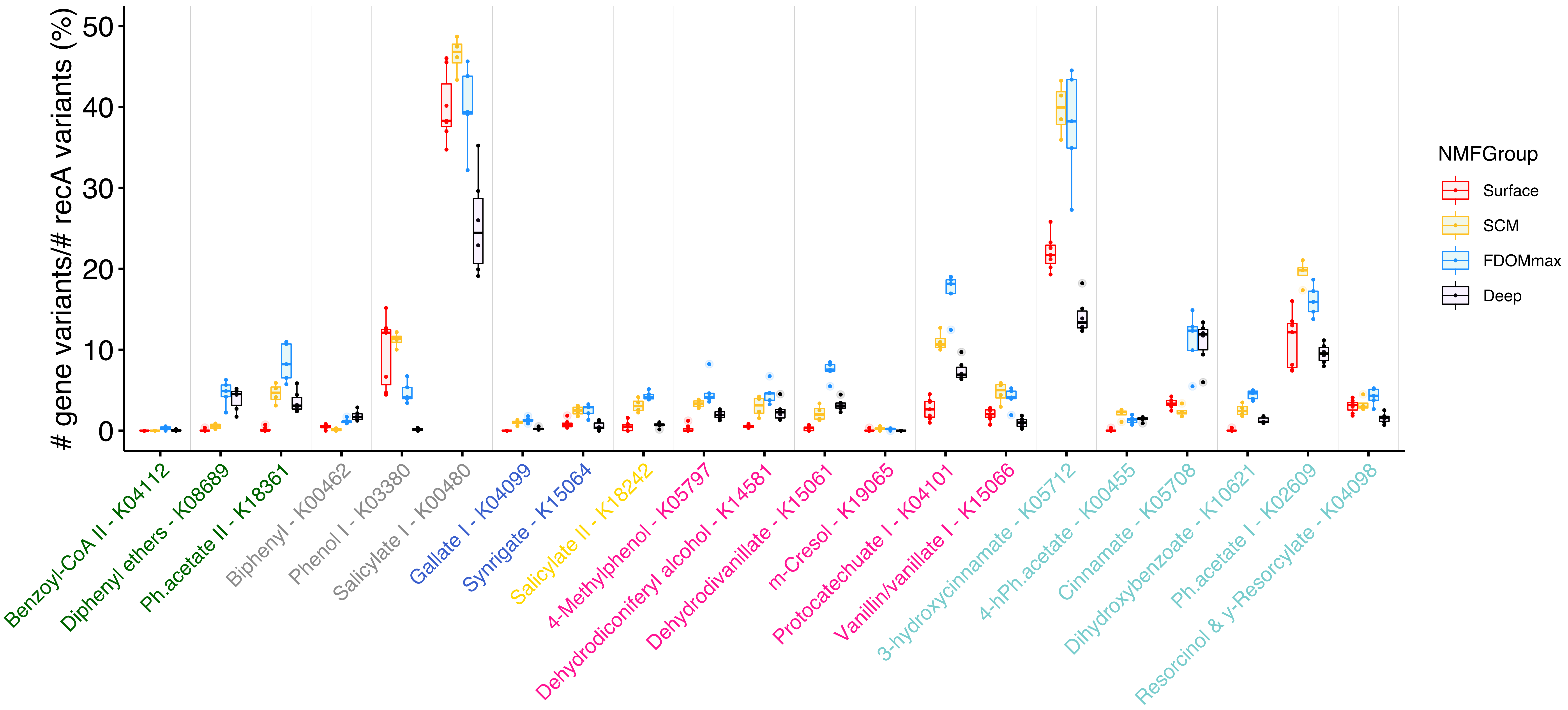

### Figure S8

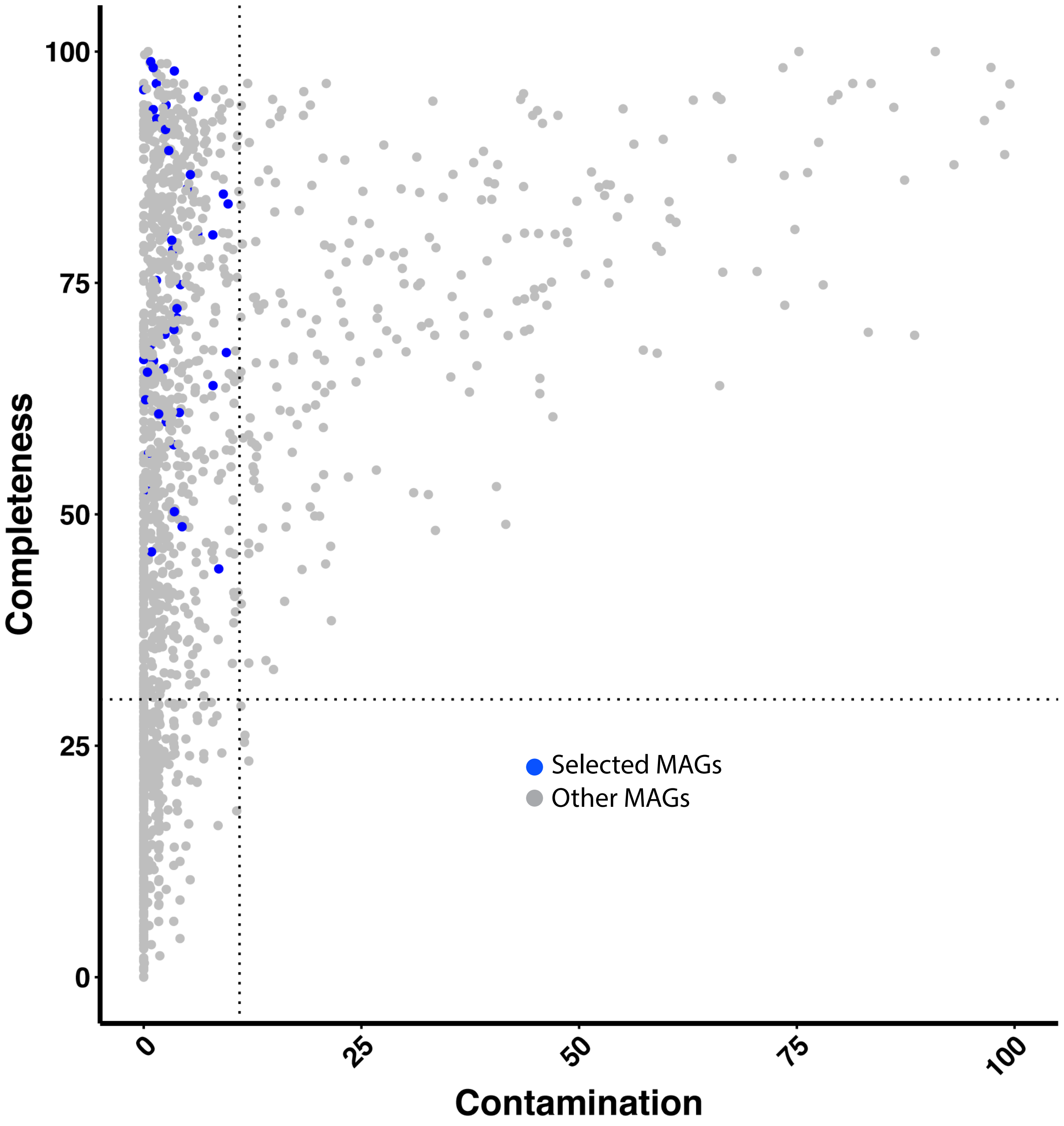

### Figure S9

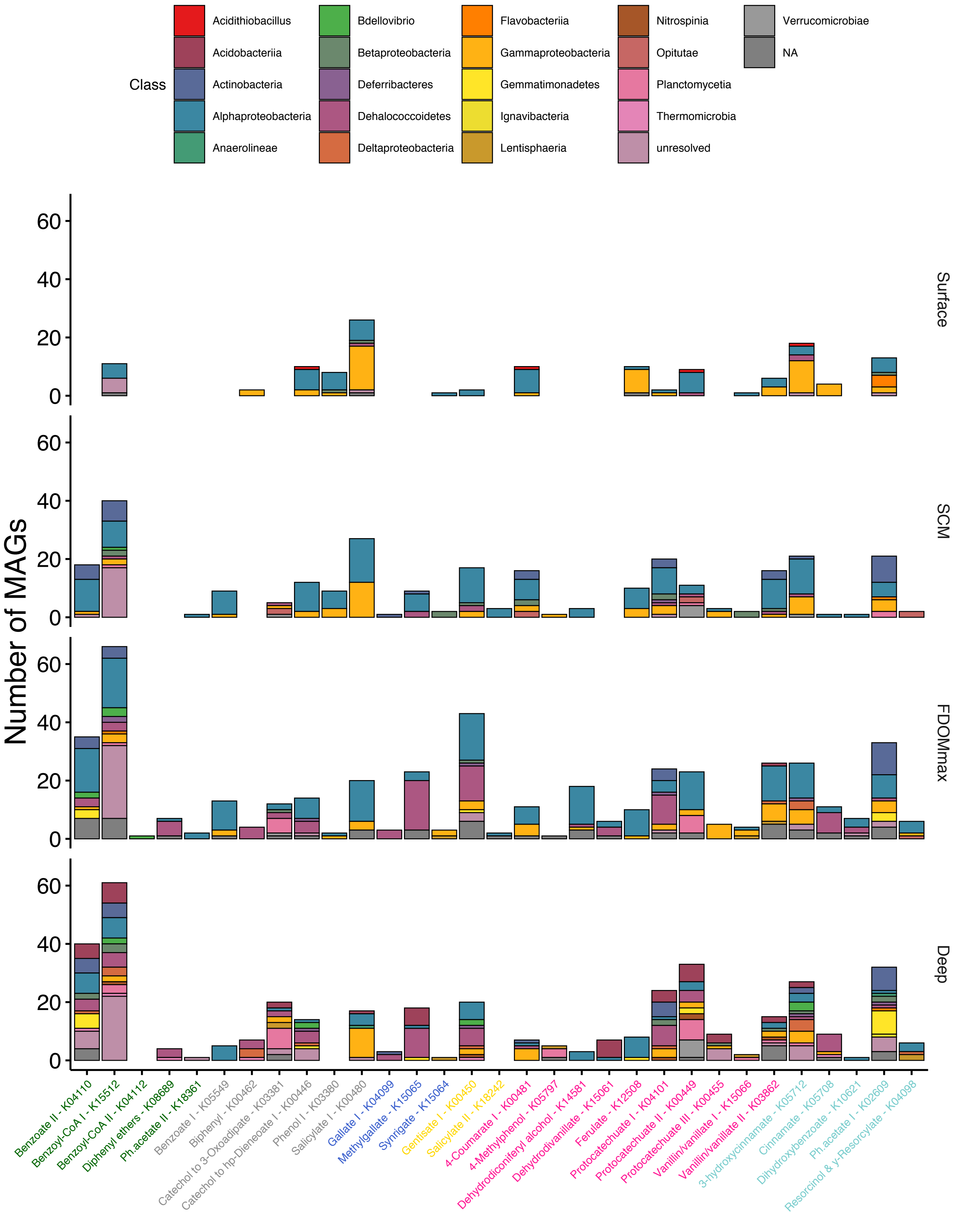

### Figure S10

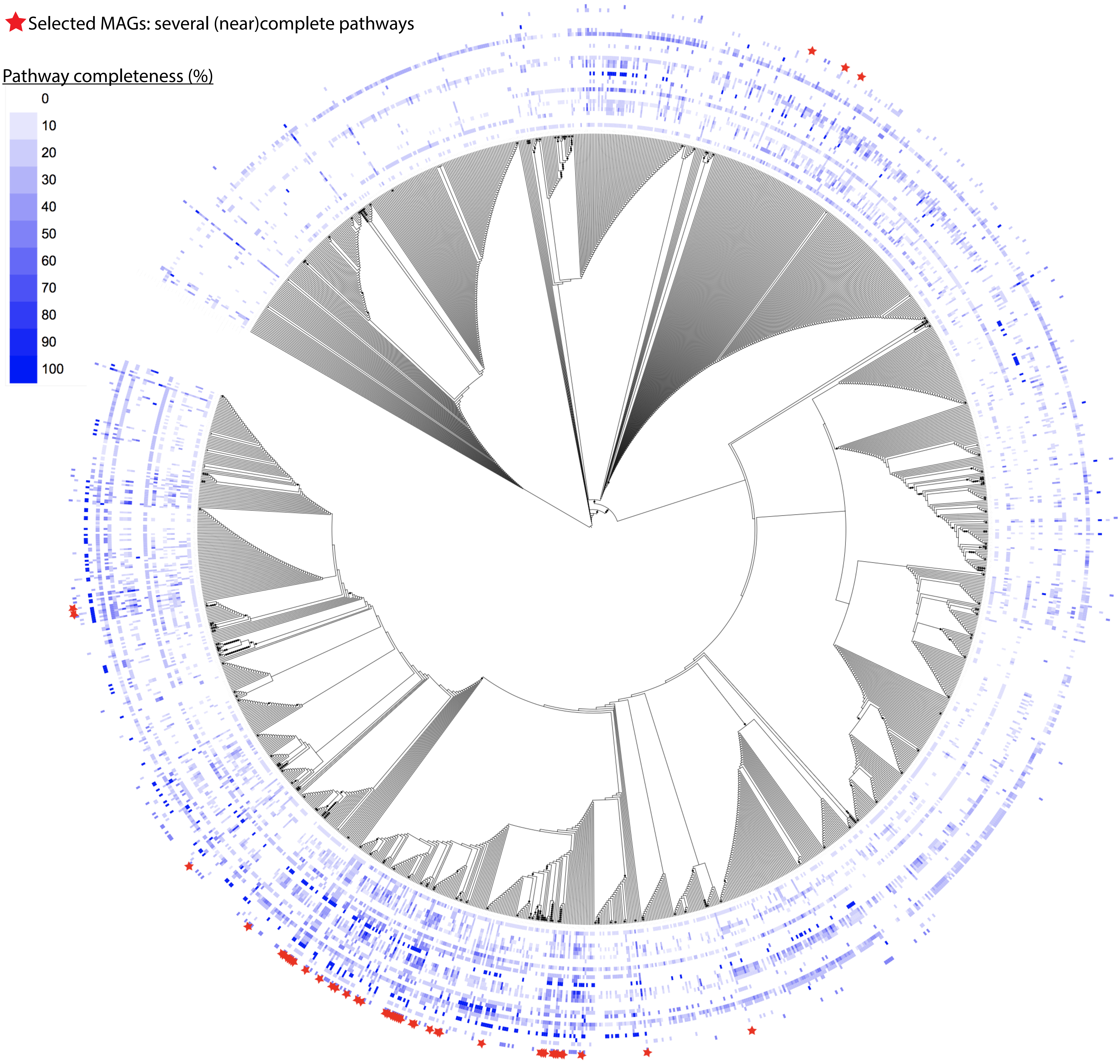

### Figure S11

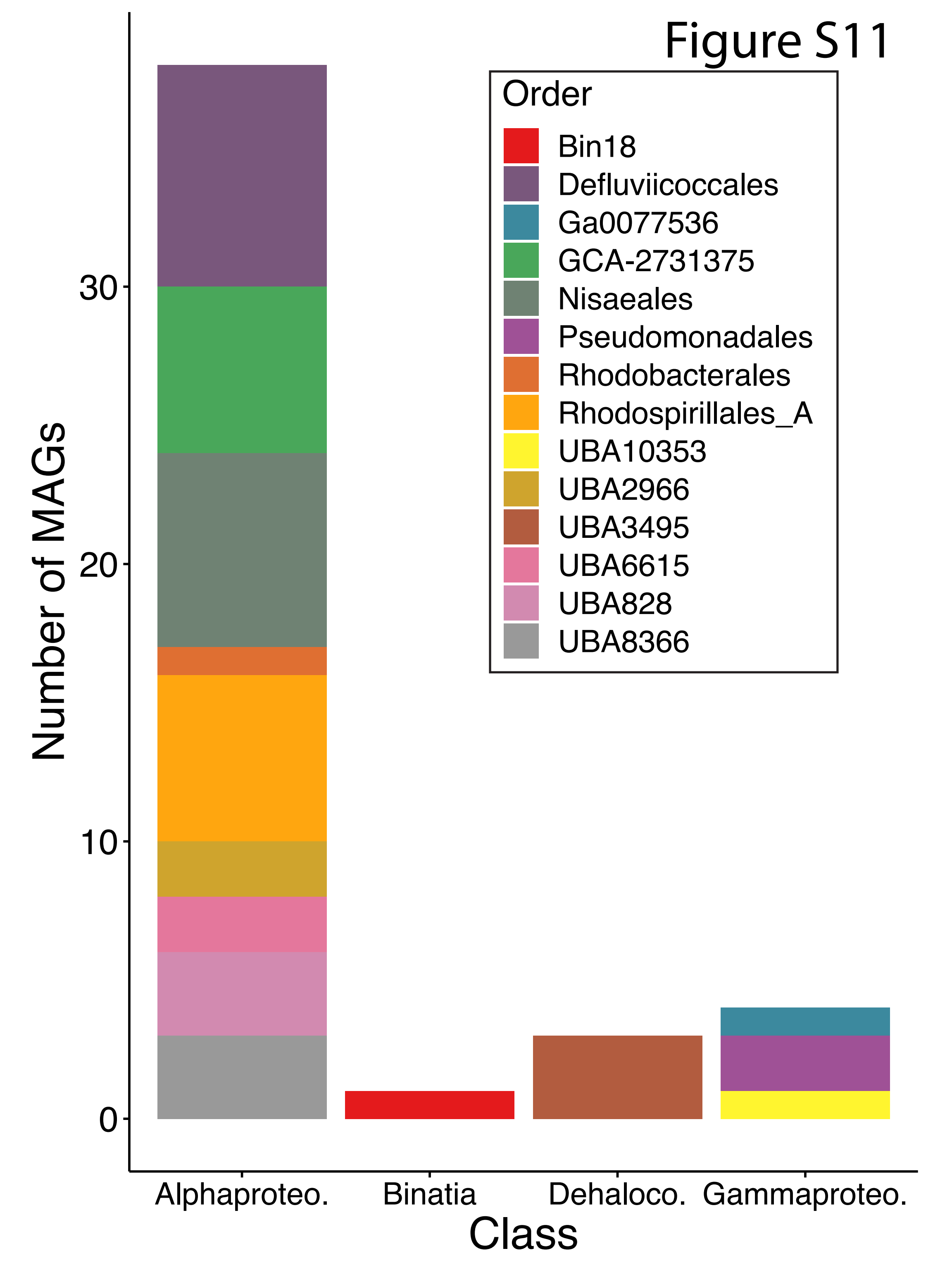

### Figure S12

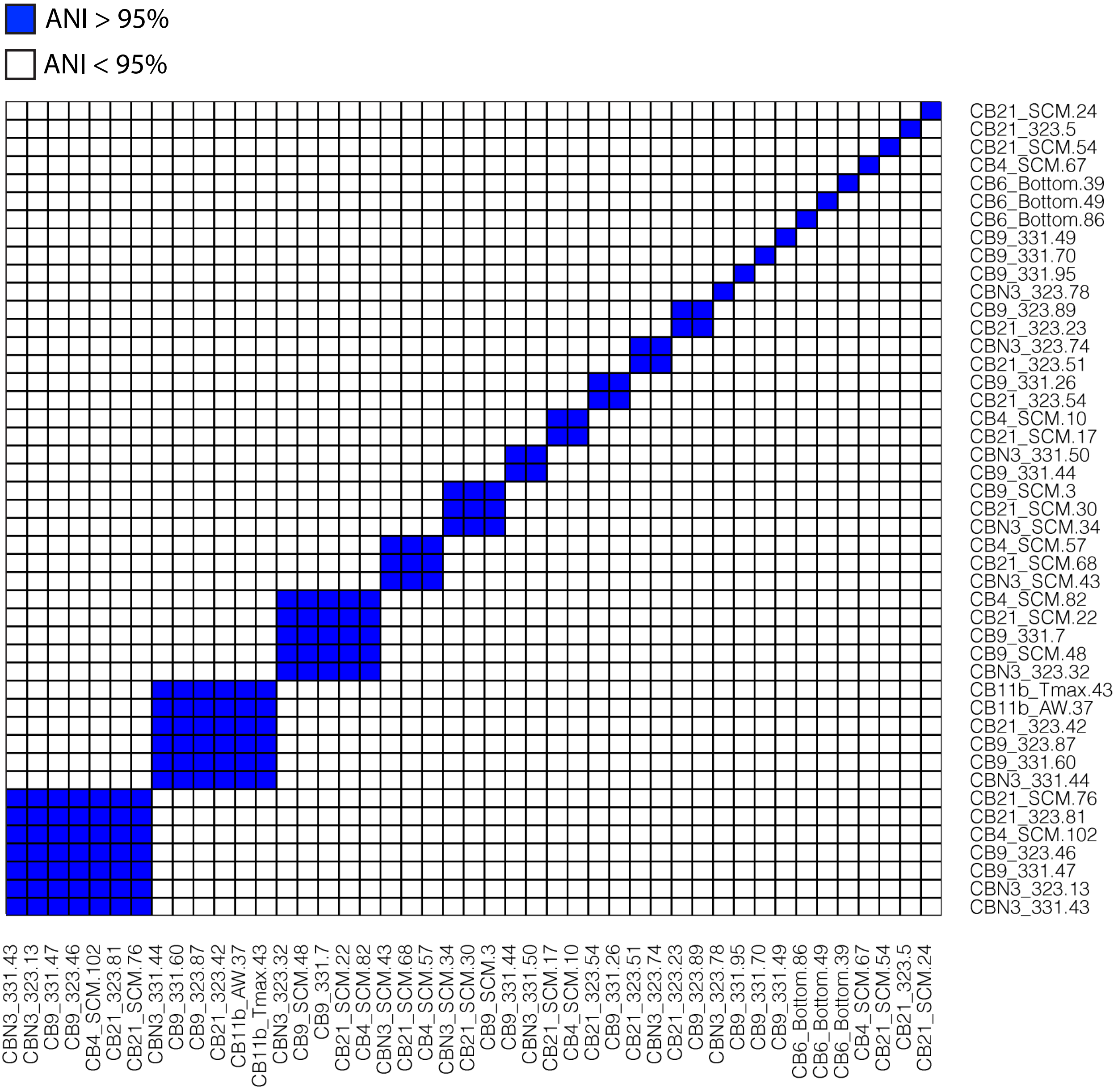
